# Rapid SARS-CoV-2 Adaptation to Available Cellular Proteases

**DOI:** 10.1101/2020.08.10.241414

**Authors:** M. Zeeshan Chaudhry, Kathrin Eschke, Markus Hoffmann, Martina Grashoff, Leila Abassi, Yeonsu Kim, Linda Brunotte, Stephan Ludwig, Andrea Kröger, Frank Klawonn, Stefan Pöhlmann, Luka Cicin-Sain

## Abstract

Since the pandemic spread of SARS-CoV-2, the virus has exhibited remarkable genome stability, but recent emergence of novel variants show virus evolution potential. Here we show that SARS-CoV-2 rapidly adapts to Vero E6 cells that leads to loss of furin cleavage motif in spike protein. The adaptation is achieved by asymptotic expansion of minor virus subpopulations to dominant genotype, but wildtype sequence is maintained at low percentage in the virus swarm, and mediate reverse adaptation once the virus is passaged on human lung cells. The Vero E6-adapted virus show defected cell entry in human lung cells and the mutated spike variants cannot be processed by furin or TMPRSS2. However, the mutated S1/S2 site is cleaved by cathepsins with higher efficiency. Our data show that SARS-CoV-2 can rapidly adapt spike protein to available proteases and advocate for deep sequence surveillance to identify virus adaptation potential and novel variant emergence.

**Significance Statement:** Recently emerging SARS-CoV-2 variants B1.1.1.7 (UK), B.1.351 (South Africa) and B.1.1.248 (Brazil) harbor spike mutation and have been linked to increased virus pathogenesis. The emergence of these novel variants highlight coronavirus adaptation and evolution potential, despite the stable consensus genotype of clinical isolates. We show that subdominant variants maintained in the virus population enable the virus to rapidly adapt upon selection pressure. Although these adaptations lead to genotype change, the change is not absolute and genome with original genotype are maintained in virus swarm. Thus, our results imply that the relative stability of SARS-CoV-2 in numerous independent clinical isolates belies its potential for rapid adaptation to new conditions.

## INTRODUCTION

Coronavirus disease COVID-19 emerged from Wuhan city in December 2019 and since has killed more than 2.6 million people worldwide. The pandemic spread of the disease that affected all human societies was found to be caused by a novel beta coronavirus, called severe acute respiratory syndrome coronavirus 2 (SARS-CoV-2), which is closely related to severe acute respiratory syndrome coronavirus (SARS-CoV)(1, 2). The typical, crown-like appearance of the CoVs is attributed to homotrimers of spike (S) protein (3), which mediate binding to the host cell receptors and subsequent membrane fusion. The CoV S protein can be subdivided into two functional subunits: S1 and S2 (4). In case of SARS-CoV-2, the S1 subunit directly binds to angiotensin-converting enzyme 2 (ACE2), which has been identified as the cellular receptor for SARS-CoV and SARS-CoV-2 (5, 6). Upon ACE2 binding, S needs to be primed by cellular proteases that leads to cleavage at the S1/S2 junction and the S2’ site (5, 7). This exposes the fusion peptide in the S2 subunit, allowing the fusion of viral and cellular membranes.

CoVs can enter the cell by fusing the virus envelope with either the endosomal or the cytoplasmic membrane, depending on the localization and availability of the cellular proteases in the host cell. Hence, cellular proteases have a direct impact on the cellular tropism and pathogenesis of CoVs. Different CoVs have evolved multiple ways to achieve proteolytic priming. A wide diversity of cellular proteases, including trypsin, endosomal cathepsins, transmembrane serine proteases (e.g., TMPRSS2) and furin are known to be involved in the priming of CoV spike (7). The SARS-CoV-2-S protein harbors a multibasic cleavage site at S1/S2, as opposed to the monobasic site that is present in the SARS-CoV-S (5). The multibasic site at S1/S2 also constitutes a putative furin cleavage site (RRAR). The consensus sequence for furin cleavage site is widely described as R-X-K/R-R↓ (8, 9). The S protein of numerous betacoronaviruses can be activated by furin at S1/S2 site; here, furin typically recognizes and cleaves at RRXRR motif (7, 10). SARS-CoV and murine hepatitis virus strain 2 (MHV-2) are two prominent exceptions of the betacoronavirus family that lack a furin cleavage site and need to be cleaved by other cellular proteases (7).

Interestingly, the furin recognition sequence present in SARS-CoV-2 S is identical (however, in opposite orientation) to the putative heparan sulfate (HS) interaction consensus sequences (XBBXBX, where B is a basic amino acid) (11). It has been shown that cell culture adapted Sindbis virus can mediate attachment to cell surface HS via furin cleavage site (8). Similarly, upon *in vitro* culturing, CoVs often exhibit a tradeoff between the furin cleavability of S protein and HS binding (10, 12). Thus, some CoVs can adapt to use HS as entry receptor in cultured cells (12). Recently, SARS-CoV-2 mutants with deletions at the S1/S2 junction have been described to emerge upon culturing the virus in Vero E6 cells (13). Clinical SARS-CoV-2 isolates, identified during the course of the current pandemic, have shown remarkably few mutations in the consensus sequence (14). However, several newly emerged SARS-CoV-2 variants seem to harbor mutations in S protein and exhibit increased transmission. For instance, the SARS-CoV-2 variant B.1.1.7 that emerged in the United Kingdom has been associated with a surge of COVID-19 cases (15).

In order to understand the dynamics of SARS-CoV-2 adaptation, we serially passaged virus strains in defined cell types and controlled environment. We show that SARS-CoV-2 rapidly adapts to culture conditions, by natural selection of pre-existing subdominant population in virus swarm. Furthermore, we show that the loss of the furin cleavage motif in Vero cells is due to a selection for virus variants with S protein optimized for efficient cleavage with cathepsins. However, viruses with the intact motif are retained as subdominant population in the swarm. Therefore, re-passaging of such mutated swarms on cells expressing TMPRSS2 results in a prompt reversion to the original genotype observed in low-passage clinical isolates.

## RESULTS

### SARS-CoV-2 rapidly adapted upon culture on Vero E6 cells

We performed deep sequencing of minimally passaged SARS-CoV-2 isolates to investigate the genome diversity and presence of minor variants. We observed considerable diversity in Ischgl (NK) and locally isolated Braunschweig (Br) strain that have been passage in vitro for four and 2 passages, respectively (Figure 1A-B). NK-P4 showed around 12 thousand nucleotide positions (12,406) with minor variants at more than 0.1% frequency and 1244 positions with >1% mutation frequency. Similarly, Br-P2 had 13,211 and 1175 positions with >0.1% and >1% mutation frequency, respectively. In order to exclude that the observed diversity is due to in vitro passaging, we also analyzed publically available SARS-CoV-2 RNA-Seq data (GSA accession number CRA002390) from bronchoalveolar lavage (BAL)(16). SARS-CoV-2 sequence from two patients showed high diversity similar to our minimally passaged isolates. Patient one sequence had 17410 positions with >0.1% and 2163 with >1% mutation as compared to consensus genotype, while SARS-CoV-2 genome from patient two had 6920 positions with >0.1% mutation frequency and 1161 positions with >1% allele frequency. On the other hand, BAC derived SARS-CoV-2 clone has been shown to possess relatively low diversity and lower number of genome position with >1% mutation frequency (17). These data suggested that SARS-CoV-2 virus quasispecies diversity despite stable consensus genotype.

**Figure 1.**
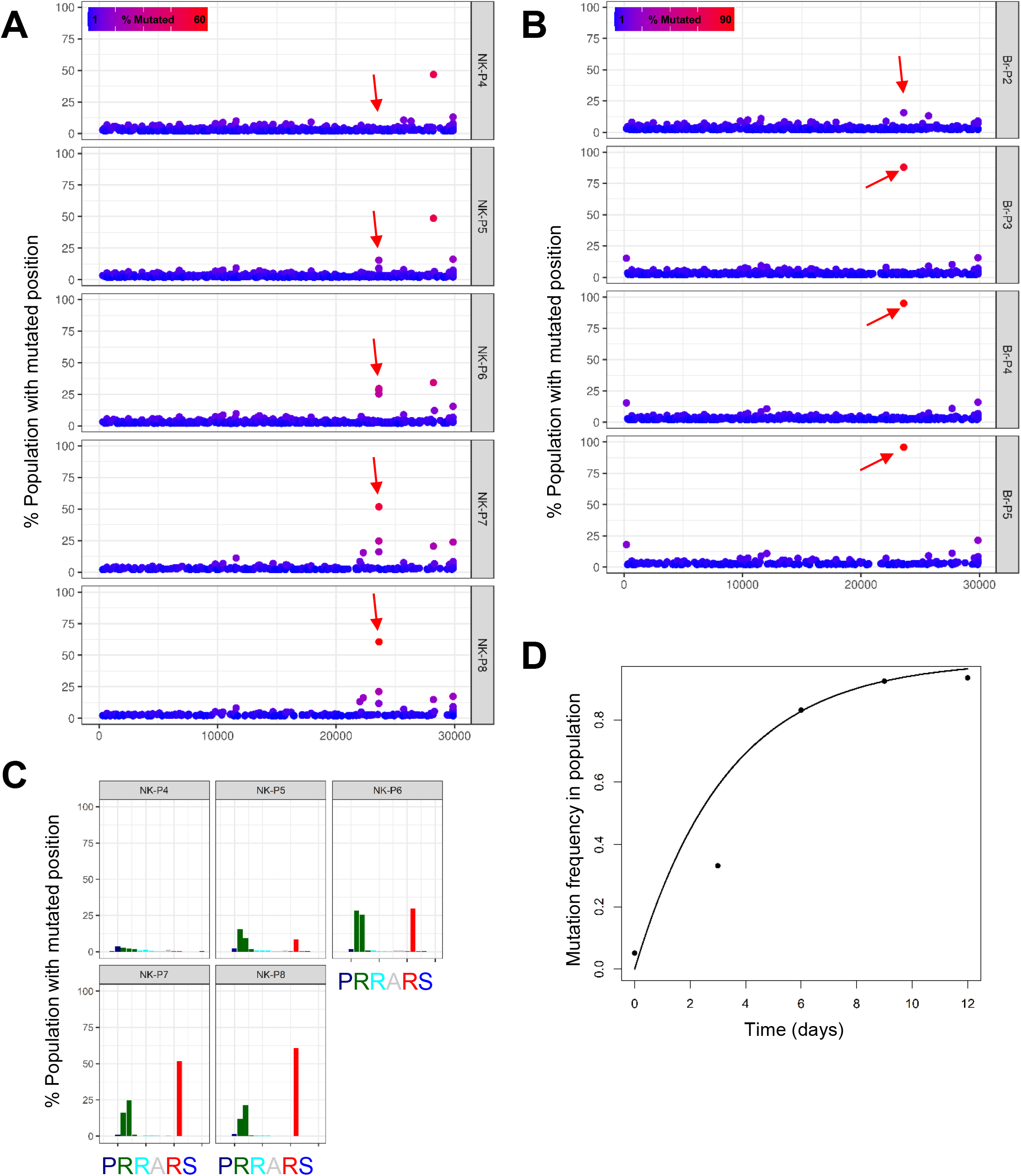
SARS-CoV-2 rapidly adapts upon passage in cultured cells. (A-B) Serial passages of Ischgl (NK) and Braunschweig (Br) strain were analyzed with deep sequencing to assess the composition of viral population upon passaging in Vero E6 cells. Each symbol represents an individual nucleotide, and genomic positions (x-axis) with mutation frequency >1% are plotted. Red arrows highlight the position of the furin cleavage site in the genome. (C) The mutation frequency of nucleotides at furin cleavage site in SARS-CoV-2 S are plotted on y-axis for NK and Br strains at different passage in Vero E6 cells. Three bars represent the nucleotide codon triplet for each corresponding amino acid at the furin cleavage site from genomic position 23,603 to 23,620 (x-axis). (D) The sum of mutation frequency at the furin site position 23,606, 23,607 and 23,616 of NK strain serially passaged on Vero E6 cells is plotted against time. The genomic positions correspond to Wuhan-Hu-1 isolate (GenBank accession no: NC_045512).

Sequence analysis of different SARS-CoV-2 clinical isolates passaged on Vero E6 cells revealed point mutation at virus genome position 23,607, which results in the loss of the furin cleavage site PRRARS at S1/S2 junction region (Table S1). These observations are in line with previous publications (13, 18). To understand how fast such mutations occur, we sequenced genomes of SARS-CoV-2 clinical isolates that were serially passaged on Vero E6 cells. Overall, the SARS-CoV-2 genomes were stable after multiple passages. However, we observed a very rapid onset of mutations in the furin recognition motif at S1/S2 site upon passaging in Vero E6 cells (Figure 1A-B). The Ischgl (NK) strain developed multiple point mutations at the furin cleavage site, resulting in loss of arginine in S protein position 682 and 685 (Figure 1A, 1C). NK strain also showed emergence of other mutation in spike with low frequency of mutated population, these are N148K (11.59%) and H245R (16.26%) in NK-P8 after eight passages in Vero E6 cells (Figure 1A). The mutation frequency for N148K and H245R increased in NK-P12 to 64.11% and 19.25% respectively (Figure S2). These S mutations were not observed in other tested strains. We used the local Braunschweig (Br) isolate to compare the dynamics of S1/S2 site mutation, as we had access to earlier passages of the Br strain. Here, a more rapid change in the composition of viral population was observed, where >85% of passage 3 (P3) genomes showed R682W mutation in S protein (Table S1; Figure S1A). We used the South Tyrol (FI) strain to compare the nature of mutation at the furin cleavage site and observed a single point mutation (R682W) in the RRAR site (Figure S1C), which was similar to Br strain. Rapid mutation in the furin site was accompanied with retention of a small fraction of genomes with wildtype furin site in all strains (Table S1). For instance, NK strain showed ~8.35% genomes with intact wildtype furin site after 12 passages in Vero E6 (Figure S2).

A mathematical model was used to fit a non-linear curve to observed mutation frequency data and calculate the rate of mutation frequency change at the furin cleavage site (Figure 1D). The NK strain showed presence of multiple sub-species including three major variants with mutation at position 23,606, 23,607 and 23,616 (Figure S1D). Thus, we summed the mutation frequencies of these three positions to model the change in mutation frequency. For NK strain, the rate was calculated as 0.3 (change in mutation frequency/day). The Br strain mutated at a much higher rate of 0.81 per day (Figure S1B). We attributes this to the high mutation frequency present in the Br strain (Br-P2).

Overall, the data demonstrated that the S1/S2 furin cleavage site was rapidly mutated upon SARS-CoV-2 culture in Vero E6 cells, but it was not completely lost and virus genomes with wildtype furin site were retained as subdominant variant in the population.

### High passage SARS-CoV-2 strains showed improved growth on Vero E6 but impaired growth on Calu-3 cells

In order to understand if the observed mutations in S influence viral growth, we compared the cell-to-cell virus spread on Vero E6 cells by assessing plaque sizes. At 3 days post infection (dpi), a significant increase in plaque sizes of high-passage virus in three independently passaged virus strains was observed (Figure 2A-B). To validate the results independently, we performed multi-step virus growth kinetics of two different virus isolates in Vero E6 cells and noticed that in both cases the high-passage virus showed significantly higher titers at 24h post infection (Figure 2C). We considered the possibility that the high-passage virus could be more efficiently released from infected cells, and compared virus titers in infected cells lysates (Figure 2D). In both conditions, the high-passage viruses showed higher titers early upon infection. Therefore, we concluded that the growth advantage of high-passage virus was a result of an early event that occurred before virus release.

**Figure 2.**
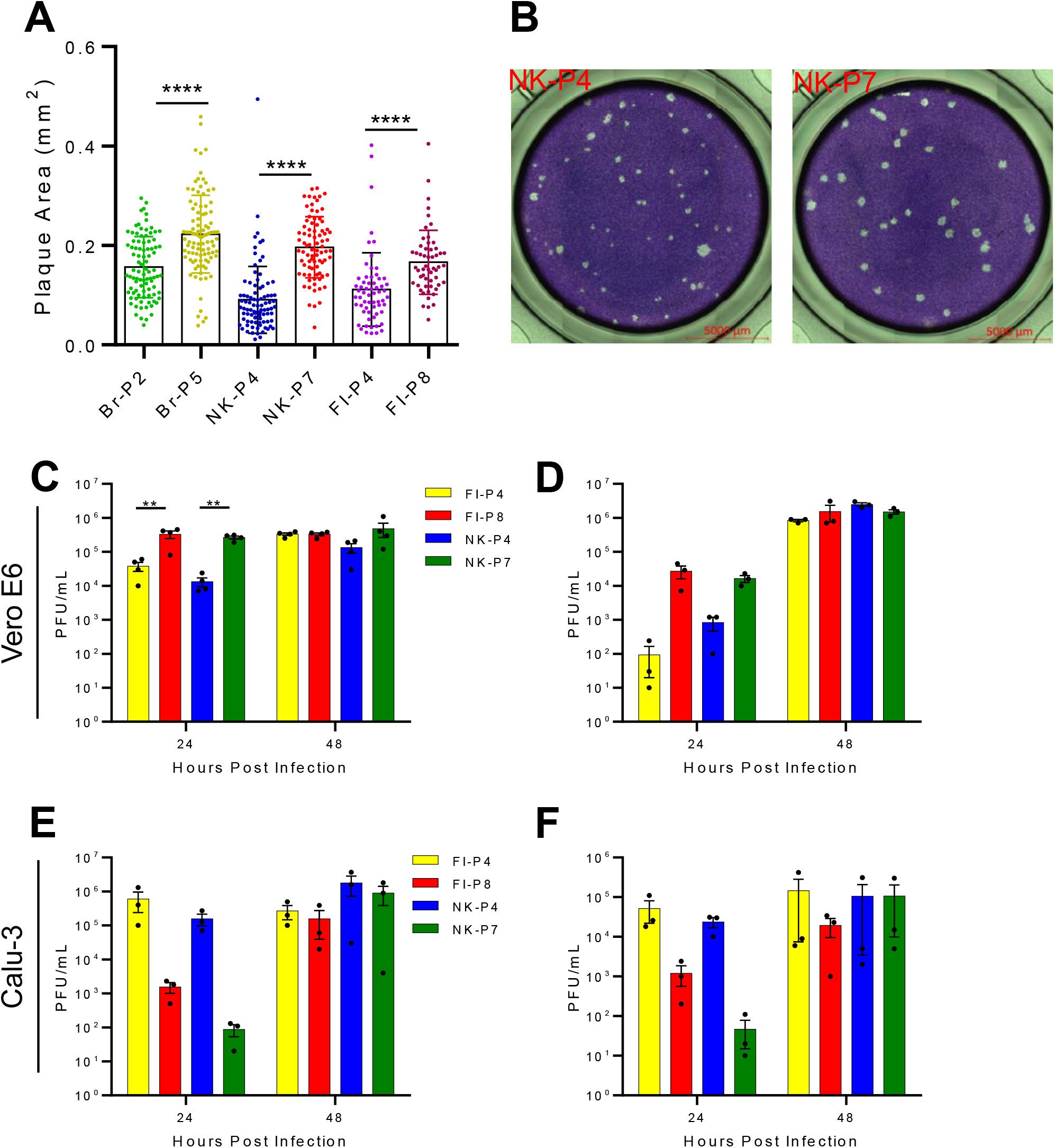
Growth properties of passaged SARS-CoV-2 strains in different cell lines. (A-B) Vero E6 cells were infected with different passages of SARS-CoV-2 strains and the plaque size was measured 3 dpi. Panel A shows the area of virus plaque on Vero E6 cells, and panel B shows representative images of two different wells infected with NK-P4 and NK-P7. (C-D) Virus growth kinetics on Vero E6 cells were established by infecting the cells at an MOI of 0.001. Supernatants (panel C) and cell lysates (panel D) were collected at indicated time points and titrated on Vero E6 cells. (E-F) Calu-3 cells were infected at an MOI of 0.001. Supernatants (panel E) and cell lysates (panel F) were collected at indicated time post infection and titrated on Vero E6 cells. Data are representative of two independent experiments. Mean and ±SEM are plotted. Each symbol in panel A represent one plaque and data are pooled from multiple infected wells of two independent experiments. Panel C-F symbols represent biological replicates. Statistical significance was calculated using one way ANOVA with Bonferroni posttest. **p < 0.01, ****p < 0.0001.

To ascertain the growth properties of Vero-passaged virus on cells expressing TMPRSS2, we tested the growth of virus strains on Calu-3 cells, and observed a mirror image of growth properties observed in Vero E6 cells. The low passage viruses grew substantially better at early time points post infection and the effects were observable equally in cell lysates and in the supernatants (Figure 2E-F). These results agree with the previous report that a multibasic cleavage site in the S protein is necessary for priming by TMPRSS2 (19).

### SARS-CoV-2 rapidly restored the furin cleavage site upon passage on TMPRSS2^+^ cells

The low passage virus growth in Calu-3 cells suggested that the furin cleavage site confers growth advantage in these cells and may result in selection of SARS-CoV-2 genomes with wildtype furin site. Therefore, we took the NK strain that had been passaged 6 times in Vero E6 cells (NK6), and re-passaged it on Calu-3 cells. SARS-CoV-2 genomes with wildtype furin site were present as a subdominant fraction (~15%) in the NK6 viral population. Upon four passages in Calu-3 cells, we observed a substantial reversion to the wildtype sequence with the intact furin site (>97%) (Figure 3A). A similar reversion was observed upon passaging the virus on Caco-2 cells: the sequence changed to a functional furin cleavage site in the vast majority of viral genomes (>95%). While high-passage viruses showed growth disadvantage in Calu-3 cells early upon infection, the virus titers rapidly recovered to low-passage levels (Figure 2E-F). The data suggest that high-passage viruses rapidly changed consensus genotype back to wildtype in Calu-3 cells. Thus, to understand the kinetics of the reversion, the virus was sequenced after each sequential passages on Calu-3. Interestingly, the majority of the reversion occurred already by on passage (Figure 3B), but a fraction of genomes with the mutated cleavage site was retained as subdominant population in the virus swarm. The composition of NK6 virus population was reshaped at reversion rate of 0.63 (change in mutation frequency/day)(Figure 3C). The consensus sequence of low-passage viruses was unaltered by their passaging on Calu-3 or Caco-2 cells (Figure 3D; Figure S3A). While S1/ S2 site mutant genomes were present at a low level, their frequency did not change. The rate of mutation frequency change suggested that the furin site mutations in both directions were a result of targeted adaptation, rather than drift. Furthermore, all viruses passaged on Calu-3 cells showed a highly similar and stable population composition, regardless of the underlying mutations induced by prior passaging on Vero cells. On the other hand, hitherto unreported mutations L27S and E-T30I in the envelope (E) protein occurred upon virus passaging in Calu-3 and in Caco-2 cells (Figure S3B-C; Table S2).

**Figure 3.**
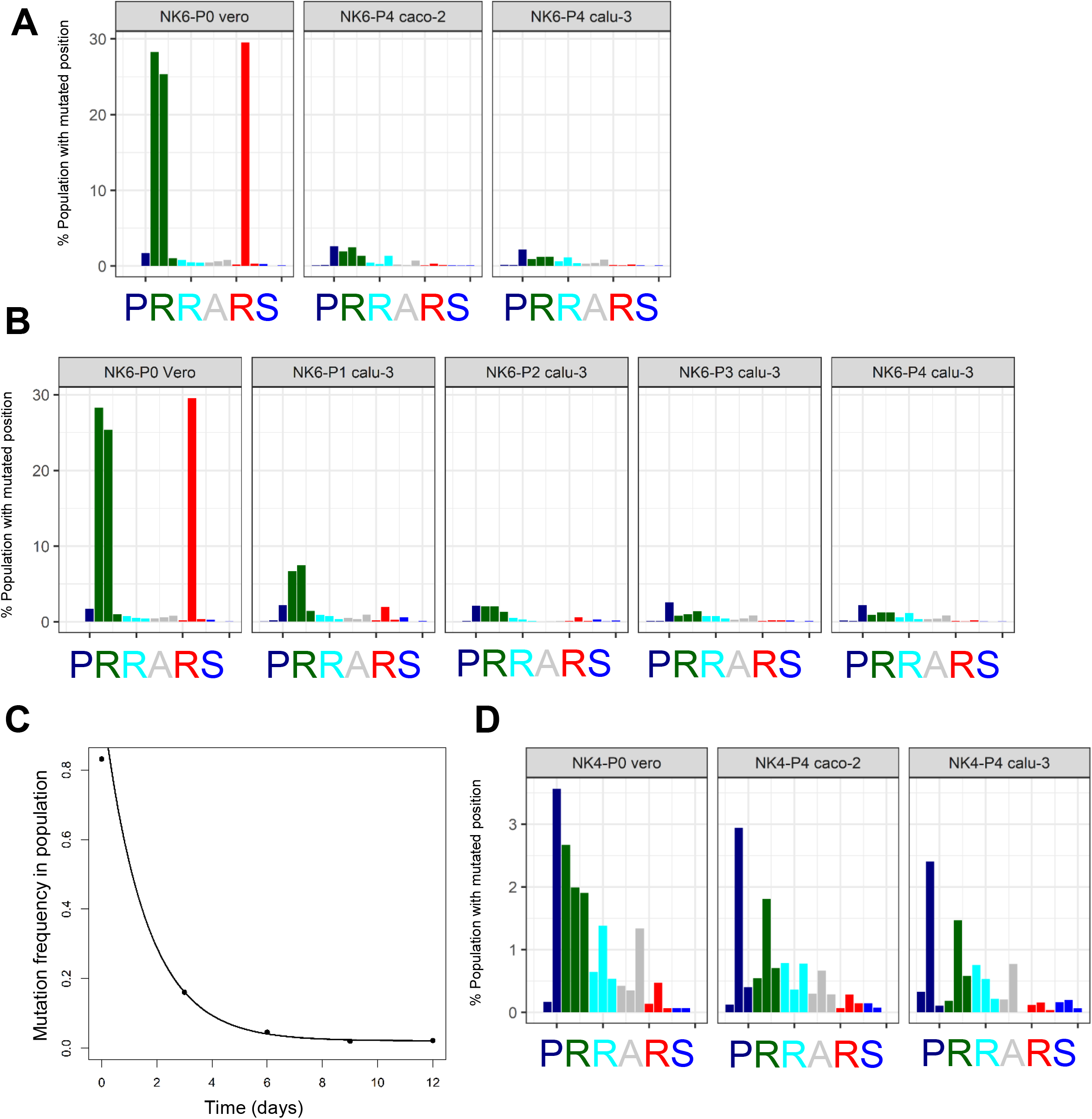
SARS-CoV-2 population composition at furin cleavage site reverts to low passage virus levels upon passage in TMPRSS2^+^ human cells. The mutation frequency of genomic position 23,603 to 23,620 (x-axis) in SARS-CoV-2 genome corresponding to amino acid codon triplets at furin cleavage site are plotted in Panel A, B and D. (A) NK strain passaged 6 time in Vero E6 cells (NK6) was serially passaged independently on Calu-3 and Caco-2 cells. P0 vero represents the genome of input virus and P4 calu-3 and P4 caco-2 are the viruses cultured on respective cells lines for 4 passages. (B) The change in SARS-CoV-2 furin site mutations is shown by plotting the nucleotide mutation frequency of NK6 virus upon serial passaging in Calu-3 cells. (C) The sum of mutation frequency at furin site position 23,606, 23,607 and 23,616 of NK6 strain serially passaged on Calu-3 cells is plotted against time. (D) NK strain passaged 4 time in Vero E6 cells (NK4) was serially passaged on Calu-3 and Caco-2 cells. P0 vero represents the genome of low passage input virus and P4 calu-3 and P4 caco-2 are the viruses cultured on Calu-3 and Caco-2 cell lines for 4 serial passages.

### Role of the SARS-CoV-2-S furin cleavage site in cell entry

Cellular proteases are critical for SARS-CoV-2 cell entry and their availability affects the virus tropism (5). Therefore, to test the role of different proteases, we infected Vero E6, Calu-3 and Caco-2 cells with low and high-passage viruses in presence of a TMPRSS2 inhibitor (camostat), a furin inhibitor (FI), or a broad-spectrum inhibitor of endosomal proteases (E-64d). The growth of all virus strains was inhibited by E-64d on Vero E6 cells, but not by camostat or FI, alone or in combination (Figure 4A). The combined use of E-64d and other protease inhibitors did not depress virus titers over the values observed in cells treated with E-64d alone (Figure S4A). This suggests that endosomal proteases, but not the cell-surface ones, were crucial for SARS-CoV-2 replication in Vero E6 cells. Importantly, no difference in titer between the high and the low-passage virus was observed in the presence of E-64d, arguing that the effect of the mutation in the high-passaged virus might occur during virus entry.

**Figure 4.**
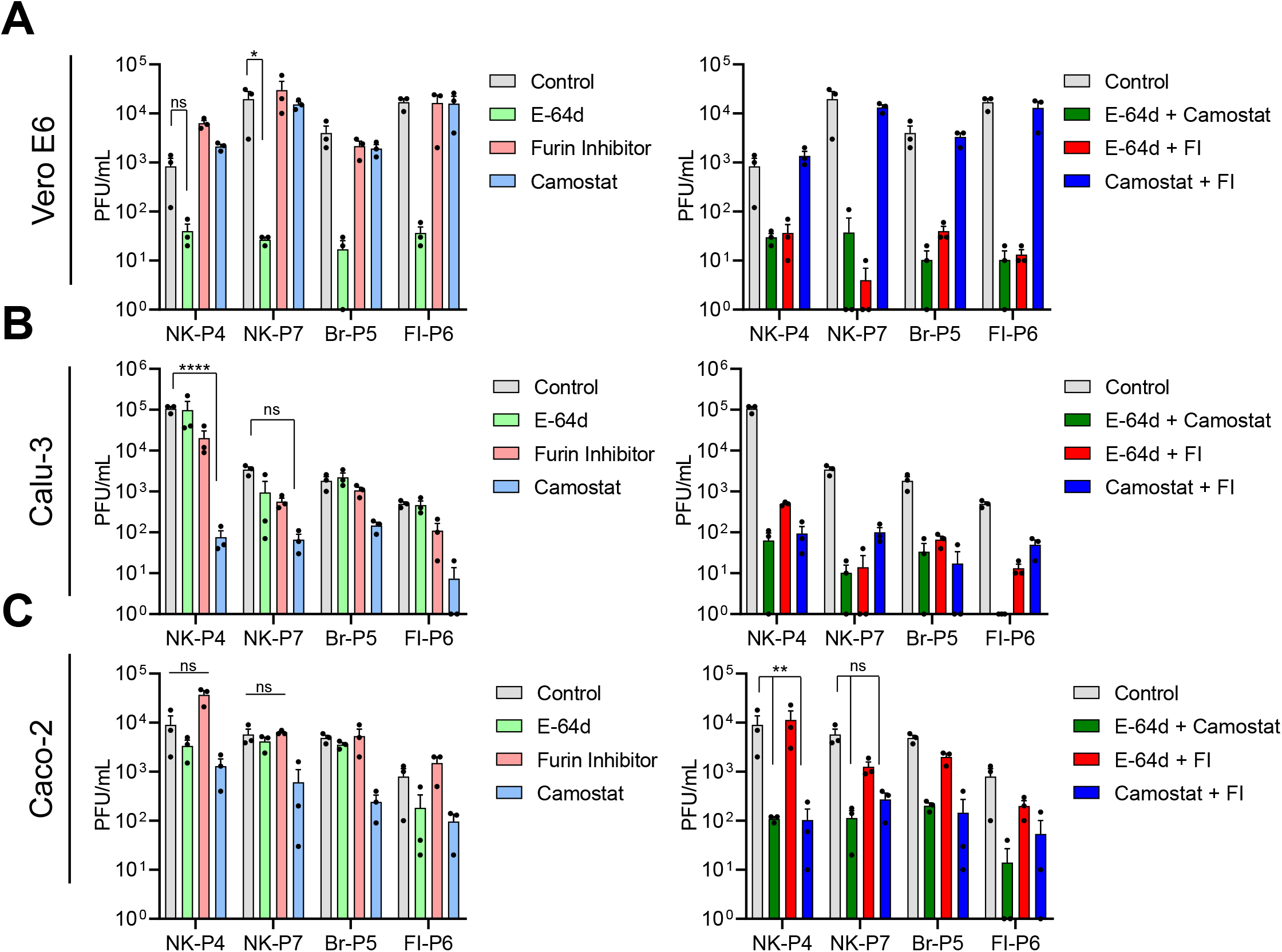
Inhibition of low and high-passage SARS-CoV-2 strain growth by different protease inhibitors. Vero E6 (A), Calu-3 (B) and Caco-2 (C) cells were infected at an MOI of 0.01 in the presence of different protease inhibitors. The supernatant from infected cells was collected at 24 hpi and titrated on Vero E6 cells. Data is representative of two independent experiments and error bars represent ± SEM of three biological replicates. Statistical significance was calculated using two-way ANOVA with Dunnett posttest, where untreated control cells served as reference. ns– p>0.05, *p < 0.05, **p < 0.01, ****p > 0.0001.

An inverse phenotype was observed in Calu-3 cells, where the TMPRSS2 inhibitor reduced the growth of all tested viruses, whereas E-64d and FI showed a less pronounced effect (Figure 4B). The differences in virus titers, seen in untreated controls, were erased in the presence of camostat. Finally, virus growth in presence of all three inhibitors was entirely abrogated in Calu-3 cells (Figure S4A). Virus replication in Caco-2 cells was not affected by inhibition of one type of protease alone, but combinations of protease inhibitors resulted in reduction of virus titers (Figure 4C). Although at 48 hpi the growth of low and high-passage viruses was similar in untreated controls, the pattern of virus growth reduction in the presence of protease inhibitors was similar to 24 hpi (Figure S3B-D).

These data suggested that growth defect of high passage SARS-CoV-2 strain is likely due to reduced entry of mutated S variants in TMPRSS2^+^ cells. We generated VSV pseudotyped virus particles that harbor wildtype SARS-CoV-2-S with PRRARS sequence at S1/S2 site, and variants with P**W**RARS or PRRA**H**S S1/S2 site sequence, two major mutated S variants that were detected in majority of passaged strains. We analyzed the VSV pseudovirus entry in Vero E6, Calu-3 and Caco-2 cells to ascertain the role of furin cleavage site in S priming and cell entry process. Viruses harboring mutated SARS-CoV-2-S showed enhanced entry in Vero E6 cells, but the entry of these variants in Calu-3 and Caco-2 was substantially lower than wildtype SARS-CoV-2-S virus (Figure 5). We concluded that the growth differences of passaged SARS-CoV-2 in different cell lines is due to the presence of mutated S1/S2 site that modulate the virus entry in these cells.

**Figure 5.**
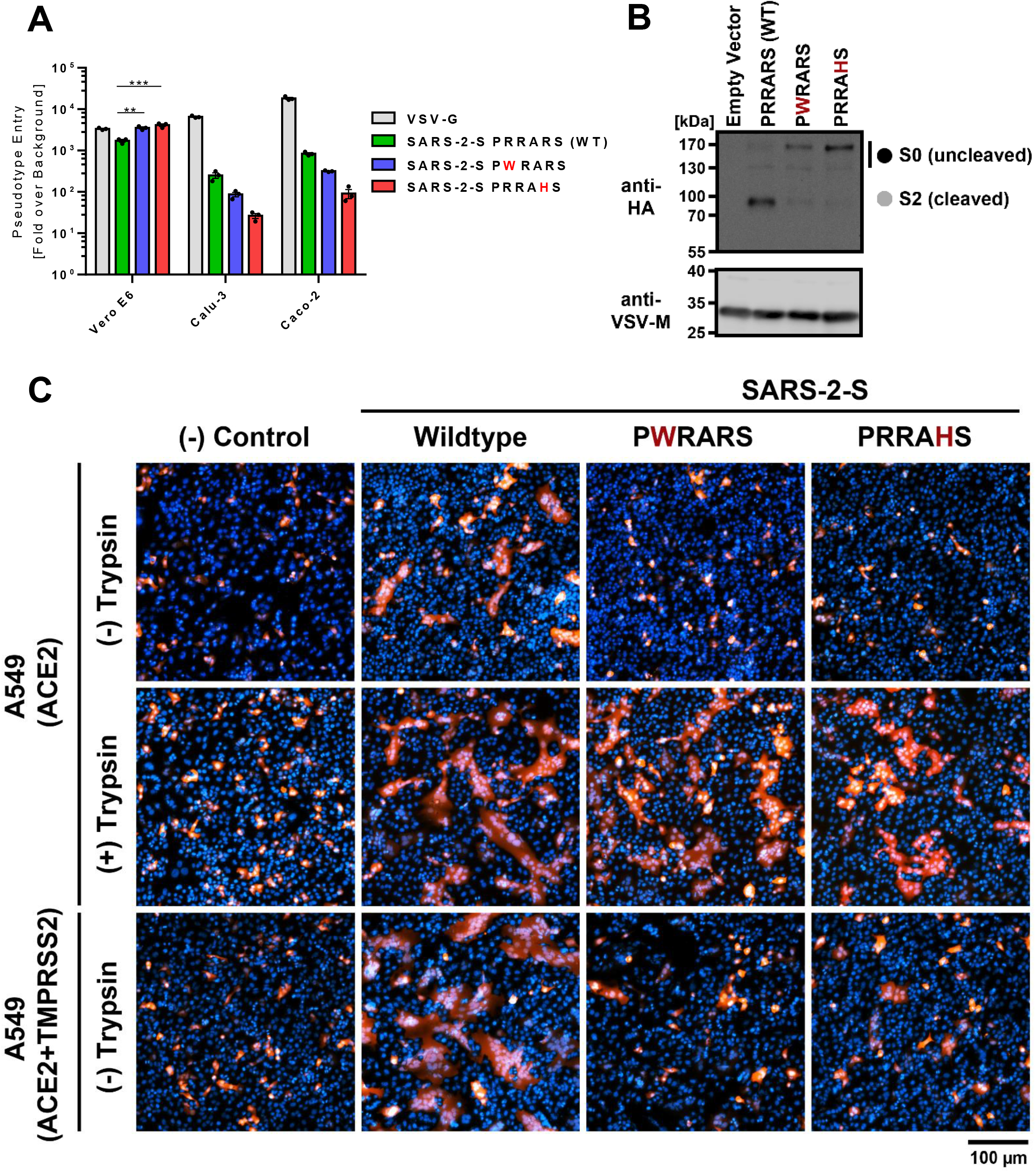
SARS-CoV-2-S furin cleavage site is required for syncytium formation and cell entry into TMPRSS2^+^ human cells. (A) Vero E6, calu-3 and caco-2 cells were infected with pseudotyped VSV harboring VSV-G, wildtype or mutated SARS-CoV-2 spike protein. At 16 h post inoculation, pseudotype entry was analyzed by determining luciferase activity in cell lysates. Signals obtained for particles bearing no envelope protein were used for normalization. The average of three independent experiments is plotted along with ± SEM. (B) (B) Analysis of furin-mediated S protein priming. Rhabdoviral particles harboring the indicated S proteins containing a C-terminal HA-tag for detection were lysed and subjected to Western blot analysis. Detection of vesicular stomatitis virus matrix protein (VSV-M) served as control. (C) Syncytium formation assay: A549-ACE2 or A549-ACE2-TMPRSS2 cells were co-transfected with DsRed expressing plasmid and vector that expressed the indicated S proteins (or no S protein, empty vector, control). At 24 h post-transfection, cells were incubated in the presence or absence of trypsin (1 μg/ml) for 1 h, before they were fixed, stained with DAPI and analyzed by florescent microscopy (scale bars, 100 μm).

### Mutated S1/S2 site is not cleaved by furin and hampers efficient cell-cell fusion

We have previously shown that PRRARS motif present at S1/S2 site of SARS-CoV-2-S matches furin consensus sequence RX[K/R]R and can be efficiently cleaved by furin, and the cleavage is inhibited with furin inhibitor (19). Here we show that mutated S proteins with PWRARS or PRRAHS sequence were not primed by furin, although wildtype variant was efficiently cleaved (Figure 5B). We next investigated if the mutated S can mediate syncytia formation in A549 cells, a cell type that has low expression of furin (20). We co-transfected cells with plasmids that expressed DsRed and S protein and observed that wildtype S protein expression resulted in multi-nucleated giant cell formation (Figure 5C), likely due to low furin expression in A549 cells. Mutated S proteins showed very few small syncytia in the absence of trypsin treatment, although trypsin treatment increase syncytia size for all conditions. Lastly, we show that TMPRSS2 expression enhanced cell-cell fusion by wildtype S protein, but the mutated S proteins could not be processed by TMPRSS2 to increase syncytia formation. The lack of syncytia formation by mutated S proteins indicated that high passage SARS-CoV-2 strains with high mutation frequency have limited cell-cell spread and likely spread via free virus particles.

### Heparan sulfate antagonist did not inhibit SARS-CoV-2 adaptation in Vero E6 cells

It has been previously reported that CoVs may trade furin cleavage sites in cell culture for a heparan sulfate (HS) binding site, thus improving virus binding to the cell surface (10, 12). However, in the case of SARS-CoV-2, the furin site PRRARS has a composition that overlaps with the HS binding motif XBBXBX. The observed mutations in HS binding motif (R682W and R685H) decreased the positive charge of the site and increased its hydrophobicity. We hypothesized that virus is mutating S1/S2 site to lose the HS binding site in the S protein, thus enhancing the growth of mutated virus in TMPRSS2^-^Vero E6. To test this hypothesis, we compared the growth of the low-passage virus to the high-passage mutants with HS antagonist in Vero E6 cells. At 24 hpi, we observed a 10-fold median increase in virus titers of the low-passage virus in the presence of HS antagonist, but the effect on high-passage viruses was non-significant (Figure 6A). HS antagonist treatment increased the virus plaque size, but the difference between low and high-passage virus remained significant (Figure 6B; Figure S5B). Furthermore, absorbing the virus on soluble or immobilized heparin resulted in a modest decrease in virus infectiousness of the low-passage virus, but no effects on the high-passage one (Figure S5A). We argued that modest HS antagonist effects might inhibit SARS-CoV-2 adaptation in the Vero E6 cells. Thus, we passaged the viruses in the presence of surfen and sequenced the genome after four passages. However, we observed that all three strains passaged in the presence of HS antagonist mutated the S1/S2 site (Figure 6C; Figure S5C-D). The data indicate that although HS antagonist modestly influence the growth of low-passage SARS-CoV-2 virus, heparan sulfate binding does not derive SARS-CoV-2 adaptation in Vero E6 cells.

**Figure 6.**
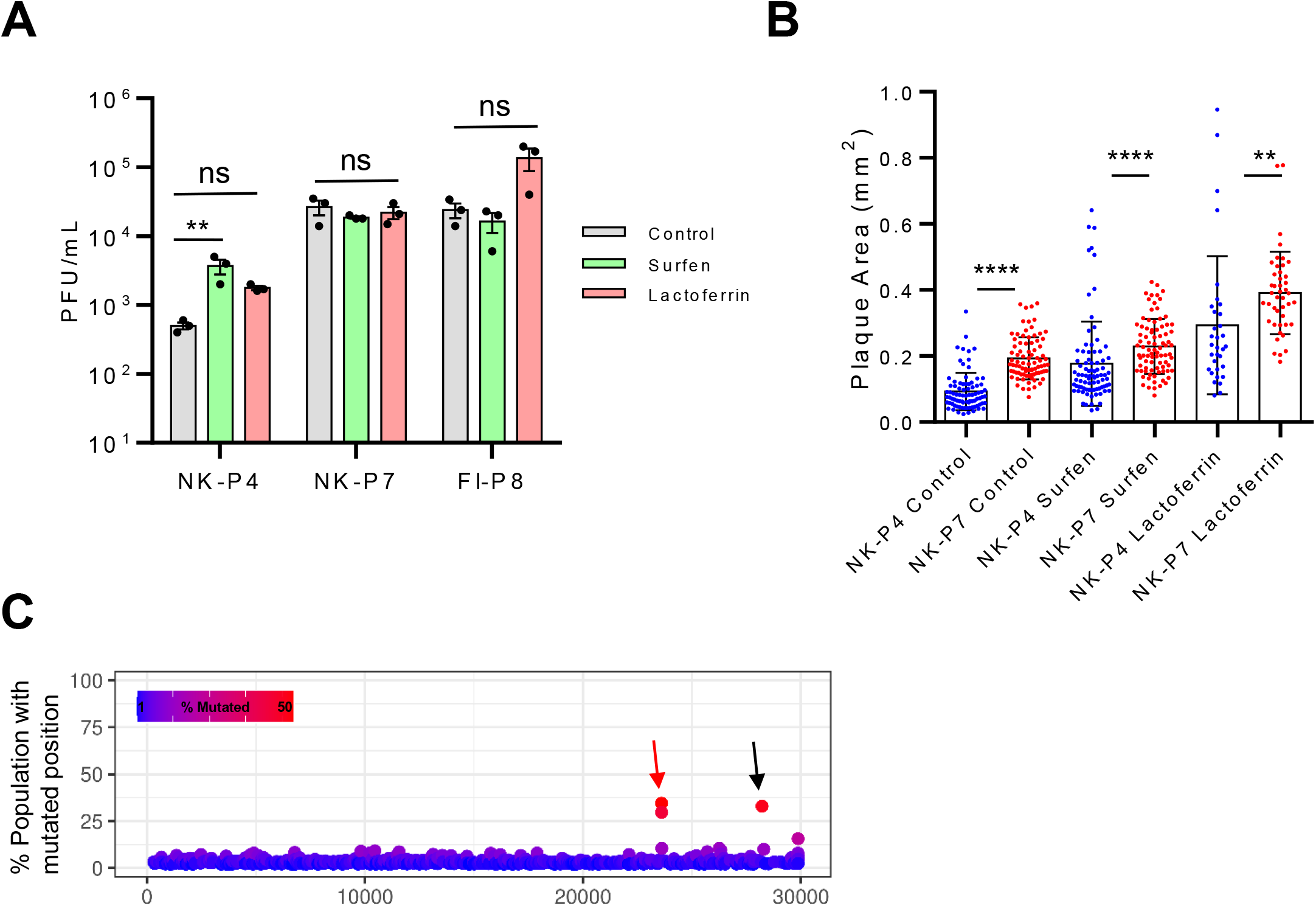
Heparan sulfate antagonists fails to inhibit SARS-CoV-2 adaptation on Vero E6 cells. (A) Vero E6 cells were infected with different SARS-CoV-2 passaged strains at an MOI of 0.001 and treated with Surfen or lactoferrin. The supernatant was collected at 24 hpi and titrated on Vero E6 cells. Data are representative of two independent experiments, and each symbol represents biological replicate. Mean and ±SEM is plotted. (B) Vero E6 cells were infected with low or high-passage NK strain viruses. The cells were overlaid with methylcellulose supplemented with 10 μM surfen or 1 mg/mL lactoferrin. The virus plaque size was quantified 3 dpi. Each symbol represents one plaque and data is pooled from multiple infected wells of two independent experiments. (C) NK strain was passaged in Vero E6 cells in the presence of 10 μM surfen. After 4 passages, virus genome sequence was analyzed with deep sequencing. Each symbol represents an individual nucleotide, and genomic positions (x-axis) with mutation frequency >1% are plotted. Red arrows highlight the position of the furin cleavage site and black arrow show synonymous mutation. Statistical significance was calculated using one way ANOVA test and Bonferroni posttest. ns – p>0.05, **p < 0.01, ****p > 0.0001.

### SARS-CoV-2 S1/S2 site adaption to available proteases

We next investigated for possible optimization of S1/S2 cleavage site in a protease specific manner. We used SARS-CoV-2-S S1/S2 site derived peptides that were fluorescently labeled with FAM-TAMRA FRET pair to assess the peptide cleavage efficiency by different proteases (Figure 7A). The wildtype SARS-CoV-2-S S1/S2 site harboring furin cleavage site was rapidly cleaved by furin, but the mutated peptides with PWRARS or PRRAHS sequence were not cleaved by furin (Figure 7B). This confirmed the VSV pseudotyped virus priming observed with western blot analysis (Figure 5B). The peptide cleavage kinetics showed that mutated peptide with PWRARS S1/S2 site was processed at a high rate by recombinant cathepsins, and PRRAHS variant was efficiently cleaved by cathepsins B. Interestingly, trypsin processed the PWRARS peptide with higher efficiency than the wildtype variant (Figure 7C). We concluded that high passage SARS-CoV-2-S is more efficiently cleaved by cathepsins, which are important for virus cell entry and S priming in Vero E6 cells. Thus, selection for SARS-CoV-2-S variant more efficiently primed by cathepsins derived virus adaptation in Vero E6 cells and lead to the emergence of S1/S2 site mutants in these cells.

**Figure 7.**
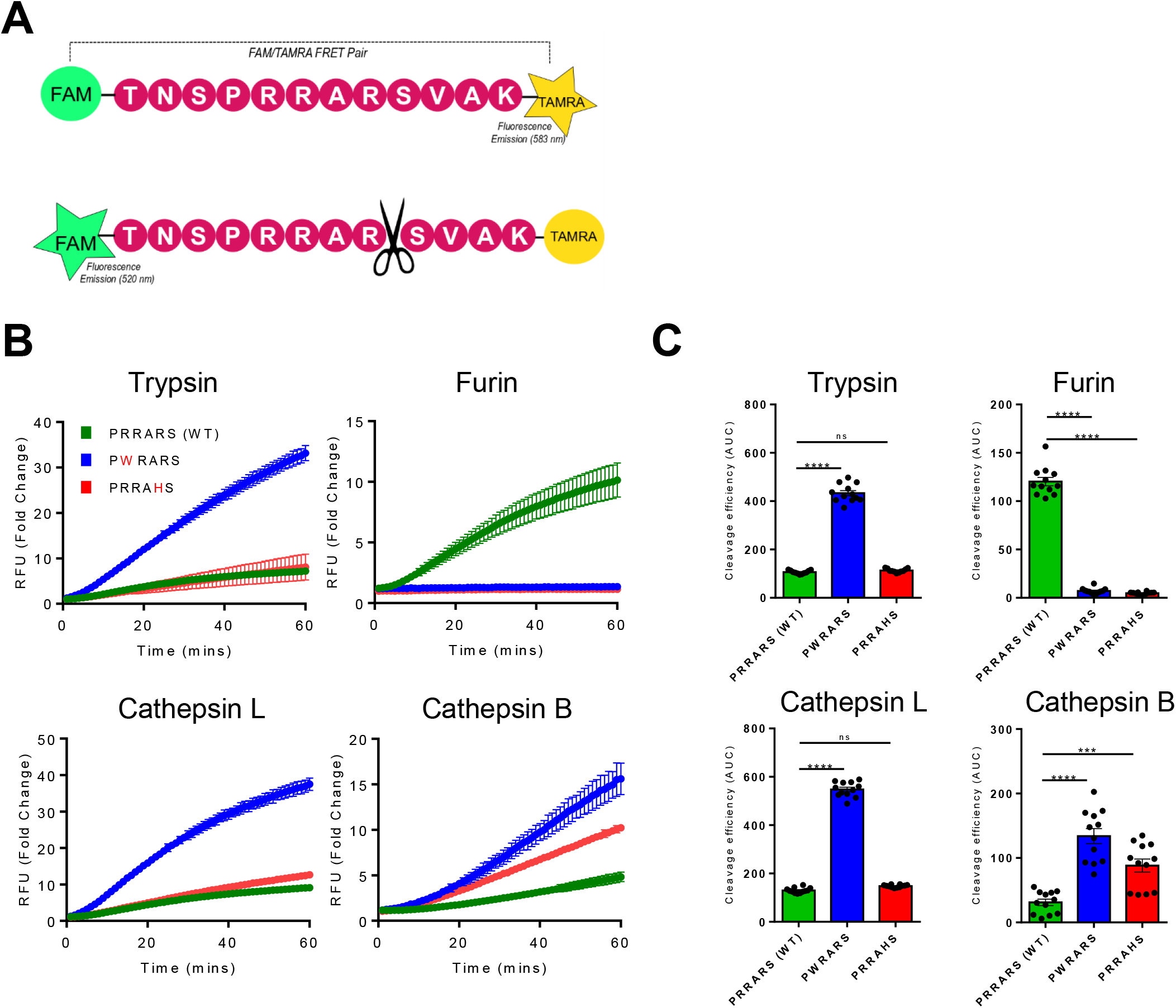
SARS-CoV-2 mutated spike variants are more efficiently cleaved by cathepsins. (A) FAM-TAMRA fret pair coupled S1/S2 spike cleavage site mimetic peptide design. (B) Fluorogenic S1/S2 spike cleavage peptides were cleaved by recombinant proteases and fold increase in FAM florescence is shown. Data is representative of 3 independent experiments, and mean ± SD of 4 replicates is plotted. (C) Peptide cleavage efficiency by different recombinant proteases was measured by calculating area under the curve (AUC) from panel B curves. Replicates from 3 different experiment are shown as mean ± SEM. Statistical significance was calculated with one way ANOVA and Bonferroni posttest. ns – p>0.05, ***p < 0.001, ****p > 0.0001.

## DISCUSSION

RNA virus populations are composed of a cloud of different genome variants known as quasispecies (21, 22) that arise due to erroneous proof-reading of RNA-dependent RNA polymerase (RdRp), and are essential for adaptive evolution and fitness of RNA viruses (23). Our deep sequencing data showed that SARS-CoV-2 exhibit remarkable genome stability overall. However, this stability does not hinder the ability of the virus to adapt quickly. Furthermore, high diversity in the clinical SARS-CoV-2 isolates as compared to BAC derived clones suggest that these minor variants play an important role in overall virus fitness and its adaptation potential (17).

Although the loss of furin site led to virus growth advantage in Vero E6 cells, genomes with wildtype motif were not completely lost. The virus population accumulated the mutations at the furin cleavage site in an asymptotic manner, stabilizing after 4-5 passages and maintaining a subdominant fraction of population with the intact motif. Mutation dynamics in Calu-3 cells followed a similar, albeit reverse, pattern and viruses with mutated PRRARS motif were retained in the virus swarm. We observed different adaptation rate in different strain that positively co-relates with the size of the subdominant population in these strains, which indicates that adaptation dynamics are likely dependent on the presence and size of subdominant populations in virus swarm. Furthermore, absence of S1/S2 mutation in BAC derived SARS-CoV-2 clones after five passages in Vero E6 cells (17) suggest that the rapid emergence of mutation that we observed is likely due to the presence of minor variants in clinical isolates. These observations suggest that SARS-CoV-2 maintains a considerable diversity in the quasispecies, facilitating natural selection and a rapid virus adaptation to changing environment and conditions. The imperfect fitness at individual level adds to the ability of the population to quickly adapt, thus contributing to the overall fitness of the swarm.

Our results are in agreement with previous report that the presence of a multibasic site is necessary for spike processing by furin and TMPRSS2 (19). In PRRAHS site, the arginine at position 685 is replaced by another basic amino acid histidine, and the ratio of hydrophilic residues change from 67% to 50%. Although PRRAHS fulfills multibasic site criteria, arginine is required at position 685 for TMPRSS2 spike priming. It is unclear if the PRRAHS site is cleaved at histidine or one of the other arginine residues.

The growth advantage on Vero E6 cells was surprising, because the furin cleavage site should have provided an advantage to viruses on cells that express furin or TMPRSS2 enzymes (Calu-3 or Caco-2), but its absence should not confer growth advantage in cells that lack these enzymes. SARS-CoV-2 furin cleavage site PRRARS has a composition that overlaps with the HS binding motif XBBXBX, and HS binding in the absence of cleavage enzymes might exert selection pressure. However, the importance of these HS binding receptors *in vivo* is debated as many viruses acquire HS binding upon cell culture adaptation (24). The SARS-CoV-2-S protein has been shown to bind HS (25) and heparin was shown to inhibit SARS-CoV-2 infection *in vitro* (26). Furthermore, Kim et al. has identified PRRARS site in the SARS-CoV-2 S protein as a putative HS binding site using an unbiased ligand-docking model (25). We report that the putative HS binding motif influence the growth of SARS-CoV-2, but virus adaptation is not affected by HS antagonists. Instead, the virus adaptation is driven by selection of S1/S2 variants that can be more efficiently cleaved by cathepsins, which are responsible for spike priming and virus entry in Vero E6 cells. The reverse adaptation to original genotype in Calu-3 cells is likely due to TMPRSS2 dependent priming, since these cells do not express robust levels of cathepsin L (27). However, the reverse adaptation in the Caco-2 cells show that priming and processing by TMPRSS2 is preferred over cathepsins.

Our study suggest that clinical SARS-CoV-2 isolates should be sequenced with deep sequencing to cover the minor genome variants present in the virus swarm instead of solely focusing on consensus genotype. The emergence of the novel SARS-CoV-2 variants highlight that virus can mutate and these novel variants show enhanced entry in human lung cells (28). Furthermore, these variants show reduced neutralization by serum from convalescent patient or vaccinated individuals. Out results argue that it is likely that these variants were present as minor subpopulations and expanded because of selection pressure. However, it is unclear if it purely to enhance virus entry in host cells or if is to evade the humoral immune response. Overall, we demonstrate a high potential of SARS-CoV-2 rapid adaptation, due to its swarm-like replication. Future research in animal models shall explore the potential for the *in vivo* adaptation of SARS-CoV-2, the quasispecies diversity and the bottlenecks imposed on the virus spread. Our research indicates that the presence of subdominant genetic variants within SARS-CoV-2 isolates needs to be considered as a potential determinant of their virulence.

## MATERIALS AND METHODS

### Cell cultures and Viruses

Vero E6 (ATCC CRL-1586), and 293T (DSMZ ACC-635) cells were maintained in DMEM medium supplemented with 10% fetal calf serum (FCS), 2 mM L-glutamine, 100 IU/mL penicillin and 100 μg/mL streptomycin. Calu-3 (ATCC HTB-55) and Caco-2 (ATCC HTB-37) were cultured in Eagle’s Minimum Essential Medium (EMEM) supplemented with 10% FCS, 2 mM L-glutamine, 100 IU/mL penicillin, 100 μg/mL streptomycin and 1x non-essential amino acid solution (Gibco MEM Non-Essential Amino Acids Solution 100X). A549 (ATCC CCL-185) cells were incubated in DMEM/F-12 medium with Nutrient Mix (ThermoFisher Scientific). All incubations of cells and virus were at 37 °C in a 5% CO_2_ atmosphere.

The SARS-CoV-2 strains used in the study are Braunschweig isolate (hCoV-19/Germany/Br-ZK-1/2020, GISAID database ID: EPI_ISL_491115), South Tyrol isolate (hCoV-19/Germany/Muenster_FI1103201/2020, GISAID database ID: EPI_ISL_463008), Ischgl isolate (hCoV-19/Germany/NK1103201/2020) and Zagreb isolate (hCoV-19/Croatia/ZG-297-20/2020, GISAID database ID: EPI_ISL_451934).

### SARS-CoV-2 passage and virus stock generation

Braunschweig isolate (Br) was derived from an oropharyngeal swab using Vero E6 cells. Zagreb isolate (Zg) was isolated in Zagreb and received as passage 2 after propagation in Vero E6 cells. Ischgl (NK) and South Tyrol (FI) strains were isolated by Stephan Ludwig lab in Muenster. Passage 2 of NK, FI and Zg strains were further propagated by passaging twice in Vero E6 cells at low MOI (multiplicity of infection) to obtain working virus stocks. For serial passaging of SARS-CoV-2 strains, Vero E6, Calu-3 and Caco-2 cells were seeded in T25 or T75 flask one day before infection. Confluent monolayers of the cells were infected with virus and virus supernatant was collected three days post infection (dpi) to further passage the virus.

Virus stocks were generated in a two-step protocol, where seeding stock was generated by infecting one T75 flask of Vero E6 cells. The seeding stock virus supernatant was collected 3 dpi and used to infect 10 T75 flasks for virus stock generation. The virus stock supernatant was collected from all flasks at 3 dpi and spun at 3000 g for 10 minutes (min) to remove cell debris. Then, the virus supernatant was concentrated using Vivaspin 20 concentrators (Sartorius Stedim Biotech) by spinning at 6000 g for 30 min. The resulting virus stock was aliquoted and stored at −80°C until further use.

### SARS-CoV-2 titration

SARS-CoV-2 was titrated with virus plaque assay on Vero E6 cells. Virus stocks were serially diluted in virus titration media (VTM, DMEM supplemented with 5% FCS, 2 mM L-glutamine, 100 IU/mL penicillin and 100 μg/mL streptomycin) and titrated on 24-well plates. The virus inoculum was added on the Vero E6 cells and incubated at 37°C. After 1 h, the inoculum was removed and the cell were overlaid with VTM supplemented with 1.5% carboxymethylcellulose (medium viscosity, C9481, Sigma-Aldrich) and incubated at 37°C. The overlay was removed from the cell at 3 dpi and the plates were fixed by submerging them in a tank of 6% formaldehyde (methanol stabilized) for at least 1 h. The cell monolayer was stained with crystal violet and the plaques were quantified by visual inspection with microscope. Virus supernatants were titrated in a similar fashion on 96-well plates.

### Growth kinetics and plaque area measurement

Vero E6, Calu-3 or Caco-2 were seeded in a well of a 96-well plate. Confluent monolayers were infected with at an MOI of 0.001. After 1 h of infection at 37°C, the inoculum was removed and replaced by normal media. Supernatants and infected cell lysates were collected at different time points post infection and virus titer were determined as described above.

To analyze cell-to-cell spread of SARS-CoV-2, the virus plaque area was determined. Vero E6 seeded in a well of a 24-well plate were infected with 30 PFU of SARS CoV 2. After 1 h of infection, the inoculum was removed and cells were overlaid with VTM supplemented with 1.5% carboxymethylcellulose (medium viscosity, C9481, Sigma-Aldrich) and incubated at 37°C. For testing the effect of HS antagonist, cells were pretreated with 10 μM surfen or 1 mg/mL lactoferrin for one hour at 37°C before infection, and infection was performed in the presence of surfen or lactoferrin. Finally, after 1 h, the virus inoculum was removed and monolayer was overlaid with VTM supplemented with 1.5% carboxymethylcellulose and 10 μM surfen or 1 mg/mL lactoferrin. Cells were fixed with 6% formaldehyde 3 dpi and stained with crystal violet. The plaque area was quantified using a Zeiss LSM 980 Microscope. The plaque area was measured with help of ZEISS ZEN lite 3.0 (Blue edition), and plotted using GraphPad Prism.

### SARS-CoV-2 infection in the presence of protease inhibitors and HS antagonists

SARS-CoV-2 cell entry inhibition was tested by seeding Vero E6, Calu-3 or Caco-2 cells in 96-well plates, one day before infection. On the day on infection, cell were treated with protease inhibitors (E-64d, 10μM; camostat, 50 10μM; furin inhibitor, 10μM) or HS antagonists (10 μM Surfen or 1 mg/mL lactoferrin) for 1 h at 37°C before virus infection. The infection was performed in the presence of protease inhibitors or HS antagonists, and after 1 h infection at 37°C, the virus inoculum was removed and normal media supplemented with inhibitors or HS antagonists was given to the cells. The virus supernatant was collected at 1 and 2 dpi and titrated on Vero E6 cells as described in SARS-CoV-2 titration section.

For analyzing the heparin binding, SARS-CoV-2 suspension containing ~1000 PFU was incubated with heparin sodium salt (100 μg/mL) for 1 h at 37°C. After incubation, the suspension was titrated on Vero E6 cells in a 24-well plate format. Similarly, for testing the ability of immobilized heparin to bind SARS-CoV-2, Heparin−biotin sodium salt was immobilized on Pierce™ Streptavidin Coated Plates (Thermo Fisher Scientific). The volume used to immobilize Heparin-biotin (100 μg/mL) was 100 μL per well. Plates were incubated at 37°C for 15 minutes and then washed three times with 1x PBS. The virus suspension (1000 PFU) was added to the heparin-coated wells along with empty streptavidin coated wells and incubated for 1 h at 37°C. Afterwards, the suspension was titrated on Vero E6 cells in a 24-well plate format.

### Pseudovirus Entry Assay

We used vesicular stomatitis virus (VSV) pseudotyped with SARS-CoV-2 S that were used according to a published protocol (29). In brief, 293T cells were transfected with expression plasmids for SARS-CoV-2 S proteins of either wildtype Wuhan/Hu-1/2019 (lineage B, with D614G mutation inserted) or variants with mutation in furin cleavage site. At 24h post transfection, the transfection medium was removed and cells were inoculated with a replication-deficient VSV vector lacking the genetic information for the VSV glycoprotein (VSV-G) and instead coding for an enhanced green fluorescent protein and firefly luciferase from independent transcription units, VSV*ΔG-FLuc (kindly provided by Gert Zimmer, Institute of Virology and Immunology, Mittelhäusern, Switzerland)(30). Following 1h of incubation at 37°C and 5% CO2, the inoculum was removed and cells were washed with PBS, before culture medium containing anti-VSV-G antibody (culture supernatant from I1-hybridoma cells; ATCC CRL-2700) was added and cells were further incubated. The pseudotype virus particles were harvested at 16-18 hpi. The culture medium was collected, centrifuged (2,000 g, 10 min, RT) to pellet cellular debris and the clarified supernatant was transferred into fresh tubes and stored at −80°C until further use. Each batch of pseudotypes was pre-tested for comparable transduction efficiencies by the respective S proteins and absence of transduction by bald control particles before being used. Furthermore, for each construct an untagged variant as well as a version containing a C345 terminal HA epitope tag was constructed.

### Western blot analysis

For the analysis of S protein processing, we subjected VSVpp harboring HA-tagged S proteins to SDS-PAGE and Western blot analysis. For this, we loaded 1 ml VSVpp onto 50 μl of a 20 % (w/v) sucrose cushion and performed high-speed centrifugation (25.000 g, 120 min, 4°C). Next, we removed 1 ml of supernatant, added 50 μl of 2x SDS-sample buffer and incubated the samples for 15 min at 96°C. Thereafter, the samples were subjected to SDS-PAGE and protein transfer to nitrocellulose membranes by Western blot. The membranes were subsequently blocked in 5 % skim milk solution (PBS containing 0.05% Tween-20 [PBS-T] and 5 % skim milk powder) for 1 h at room temperature. The blots were then incubated over night at 4 °C with primary antibody solution (all antibodies were diluted in PBS-T containing 5 % skim milk; mouse anti-HA tag [Sigma-Aldrich, H3663, 1:2,500] or VSV matrix protein [Kerafast, EB0011, 1:2,500]). Following this incubation, the blots were washed 3x with PBS-T before they were incubated for 1 h at room temperature with peroxidase-coupled goat anti-mouse antibody (Dianova, 115-035-003, 1:10,000). Finally, the blots were again washed and imaged. For this, an in house-prepared enhanced chemiluminescent solution (0.1 M Tris-HCl [pH 8.6], 250 μg/mL luminol, 1 mg/mL para-hydroxycoumaric acid, 0.3 % H2O2) and the ChemoCam imaging system along with the ChemoStar Professional software (Intas Science Imaging Instruments GmbH) were used.

### Syncytium formation assay

A549-ACE2 or A549-ACE2-TMPRSS2 cells were grown on coverslips seeded in 24-well plates and co-transfected with DsRed vector (1 μg/well) and S protein expression plasmid (1 μg/well) using Lipofectamine 2000 LTX with Plus reagent (Thermo Fisher Scientific) and OptiMEM medium (Gibco). After 6 h, the transfection solutions were aspirated and the cells further incubated for 24 h in standard culture medium. The medium was changed next day to serum free medium with or without 1 μg/ml bovine trypsin (Sigma-Aldrich) and the cells were incubated for additional 1 h at 37°C. Cells were washed with PBS and fixed with 4 % paraformaldehyde solution for 20 min at room temperature. After fixation, cells were stained with DAPI solution and analyzed by bright-field microscopy using a Zeiss LSM800 confocal laser scanning microscope and the ZEN imaging software.

### Peptide Cleavage Assay

Target cleavage peptides derived from the SARS-CoV-2 spike S1/S2 site amino acid sequence TNSPRRARSVA and flanked with FAM-TAMRA FRET pair ([FAM]TNSPRRARSVA-K[TAMRA][COOH.acetate]) were synthesized by Thermo Fisher Scientific. Wildtype peptide TNSPRRARSVA along with two mutated peptides TNSPWRARSVA and TNSPRRAHSVA harboring point mutation in amino acid sequence of furin cleavage site were synthesized. Recombinant furin was purchased from New England Biolabs. Recombinant L-1-Tosylamide-2-phenylethyl chloromethyl ketone (TPCK)-treated trypsin was obtained from Sigma-Aldrich. Recombinant cathepsin B and cathepsin L were purchased from R&D Systems.

Florescent peptides were diluted to 25 μM and the cleavage reaction was performed in 100 μL total volume. The reactions buffer was composed of 100 mM Hepes, 0.5% Triton X-100, 1 mM CaCl2 and 1 mM 2-mercaptoethanol (pH 7.5) for furin (10 U/mL); PBS for trypsin (100 ng/mL); 25 mM MES, pH 5.0 for cathepsin B (100 ng/μL); 50 mM MES, 5 mM DTT, 1 mM EDTA, 0.005% (w/v) Brij-35, pH 6.0 for cathepsin L (100 ng/μL). Reactions were performed in Varioskan Flash (Thermo Scientific) at 30 °C, and kinetic fluorescence measurements were recorded at one minute interval for 60 min with λex 495 nm and λem 520 nm wavelengths setting. Peptide cleavage efficiency was determined by calculating area under the curve (AUC) with fold increase in FAM florescence over time (time t=10 min to t=40 min).

### SARS-CoV-2 RNA isolation and sequencing

For the inactivation of SARS-CoV-2, the virus suspension was mixed 1:1 with peqGOLD TriFast (VWR). Following, RNA was extracted using the innuPREP Virus RNA Kit (Analytik Jena) according to the manufacturer’s instructions. The RNA quality was confirmed with Bioanalyzer (Agilent Technologies) and Qubit (Thermo Fisher Scientific). The sequencing libraries were prepared using NEBNext Ultra II Directional RNA Library Prep Kit (New England Biolabs) including ERCC RNA Spike-in control. The sequencing was performed with NovaSeq 6000 SP Reagent Kit (100 cycles) on NovaSeq 6000 System (Illumina).

Raw fastq files were processed to trim adaptor and low quality reads (fastp 0.20.0). The resulting sequence reads were aligned with reference (BWA 0.7.8). The alignment file was sorted (SAMtools version 1.10) and realigned (GATK IndelRealigner version 3.7). Finally, pileup file (SAMtools) was created and mutation were detected with VarScan (version 2.3.9). Mutation frequency for whole genome was calculated by extracting total read depth and reference base reads for each nucleotide from pileup file using SAMtools, where mutation frequency in the population is given as ratio of reference reads to total reads at the position. Mutation frequencies were converted to percent values and plotted with ggplot2 (R/Bioconductor 4.0). Alignment files were inspected with IGV (version 2.8.3) and Tablet (version 1.19.09.03) to access if nearby mutations in the furin cleavage site are present on same reads.

### Mathematical modelling

The mathematical model, to fit the curves for the mutation, assumes that the proportion of viruses with a mutation on the considered chromosome (SARS-CoV-2 genome positions 23606, 23607 and 23616) at time *t* depends on the initial proportion of mutated viruses *y*(0), the mutation rate *a* and the reverse mutation rate *b*. This leads to the ordinary differential equation *y*^′^ = *a*(1 − *y*) − *by* with the general solution 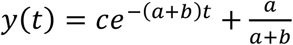. Assuming that there are no mutations at time zero, i.e. *y*(0) = 0; then 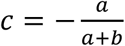 follows. If we start with a mutated strain, i.e. *y*(0) = 1, this implies 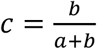. The mutation rates *a* and *b* of the corresponding non-linear curve are estimated based on the data by the optimization procedure optim provided by the R statistics software, minimizing the sum of absolute errors. R script used is provided in supplemental data.

## QUANTIFICATION AND STATISTICAL ANALYSIS

Kruskal-Wallis with Dunn’s comparison and two-way analysis of variance (ANOVA) with Dunnett posttest were used to test for statistical significance. Where appropriate, two-way ANOVA with Sidak’s multiple comparison was used. P-values < 0.05 were considered significant (*p < 0.05; **p < 0.01; ***p < 0.001), ****p < 0.0001, p > 0.05 not significant (n.s.). For all statistical analyses, the GraphPad Prism 7 software package was used (GraphPad Software).

## DATA AND CODE AVAILABILITY

The SARS-CoV-2 deep sequencing data generated during this study are available at NCBI sequence read archive. (SRA accession: PRJNA650134; https://www.ncbi.nlm.nih.gov/Traces/study/?acc=PRJNA650134). The R code used in this study are provided in supplemental data (Data S1).

## MATERIALS AVAILABILITY

Further information and requests for resources and reagents can be directed to M. Zeeshan Chaudhry and Luka Cicin-Sain. The materials will be made available upon receipt of a material transfer agreement (MTA).

## Acknowledgements

We thank HZI Genome Analytics team for support, especially Michael Jarek. We kindly acknowledge Susanne Talay and Markus Hoffmann for helpful discussion. We thank Ayse Barut and Inge Hollatz-Rangosch for technical assistance.

---

**Figure S1.**
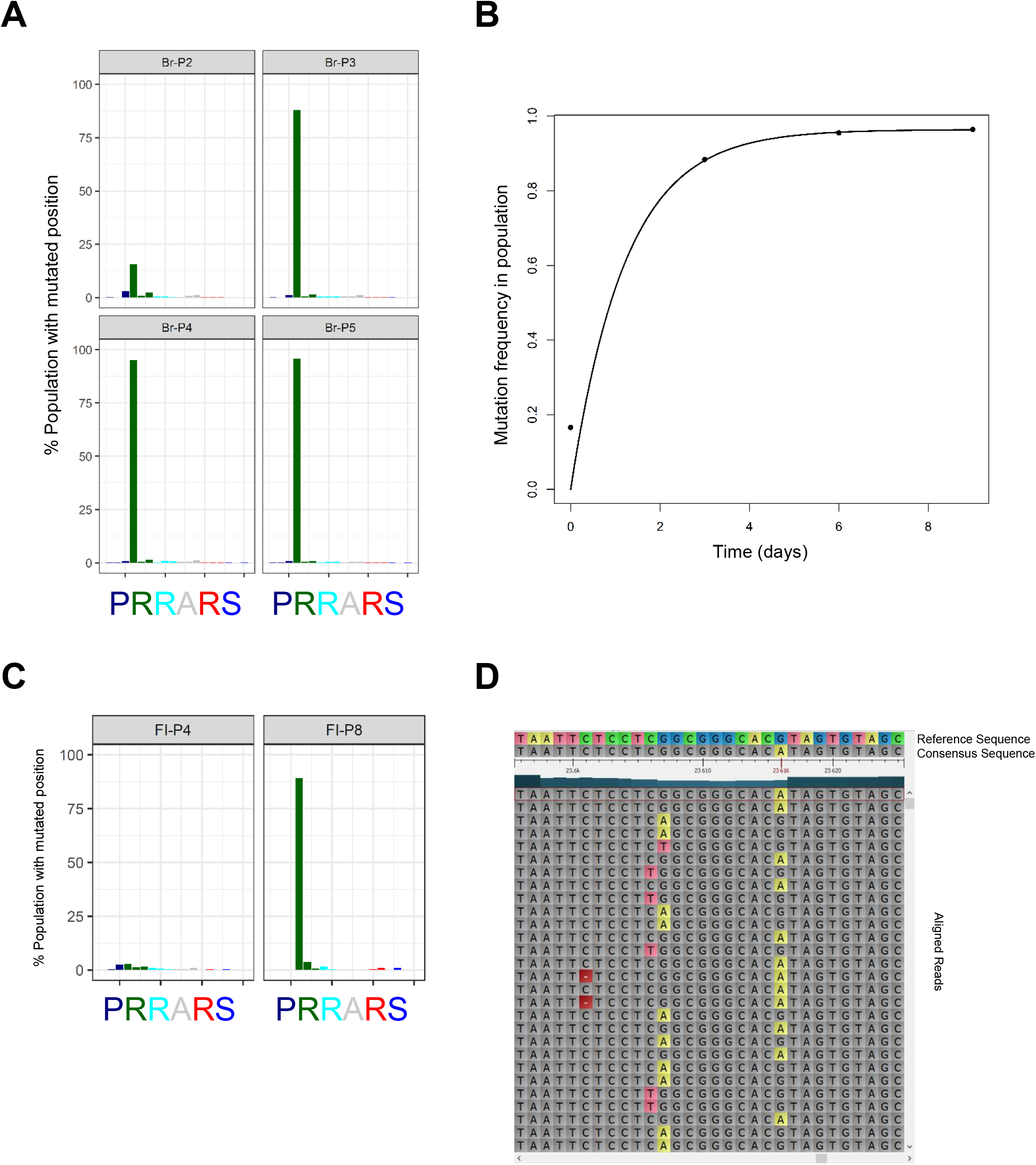

**Figure S2.**
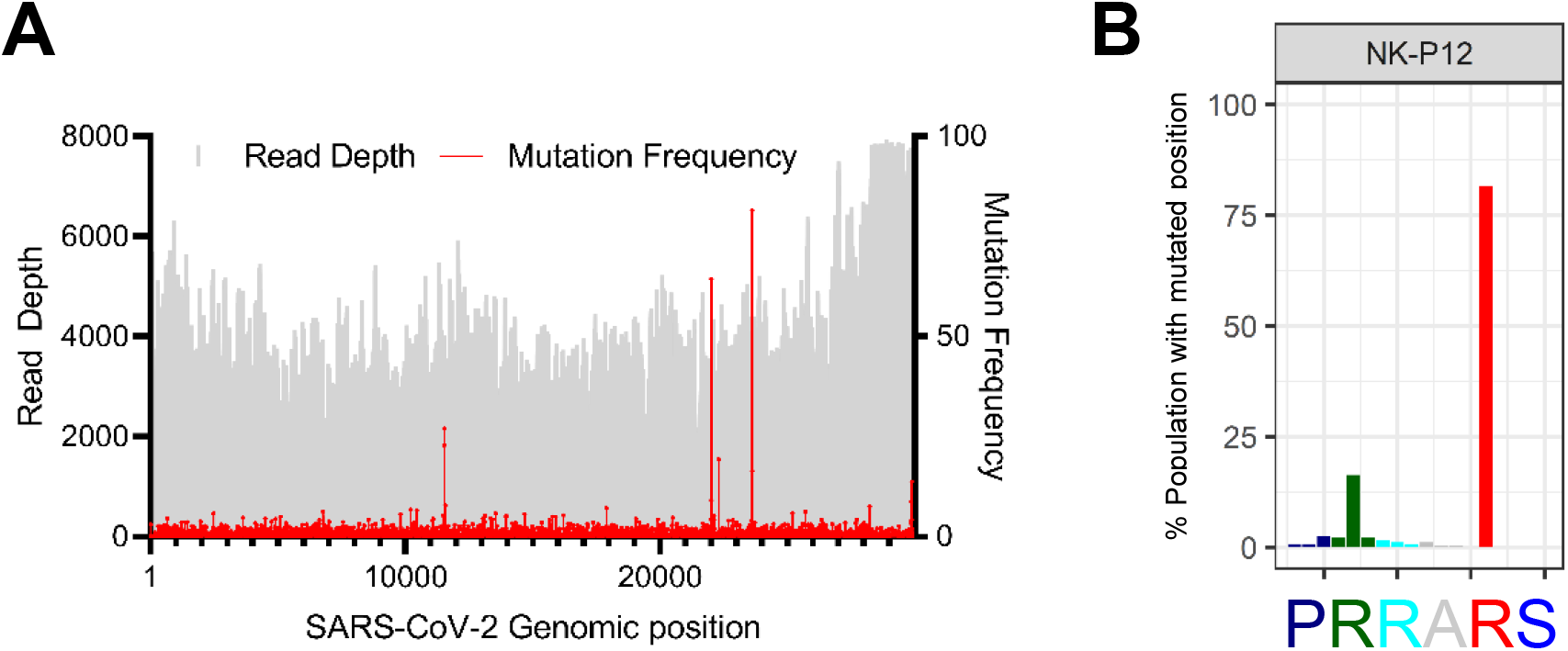

**Figure S3.**
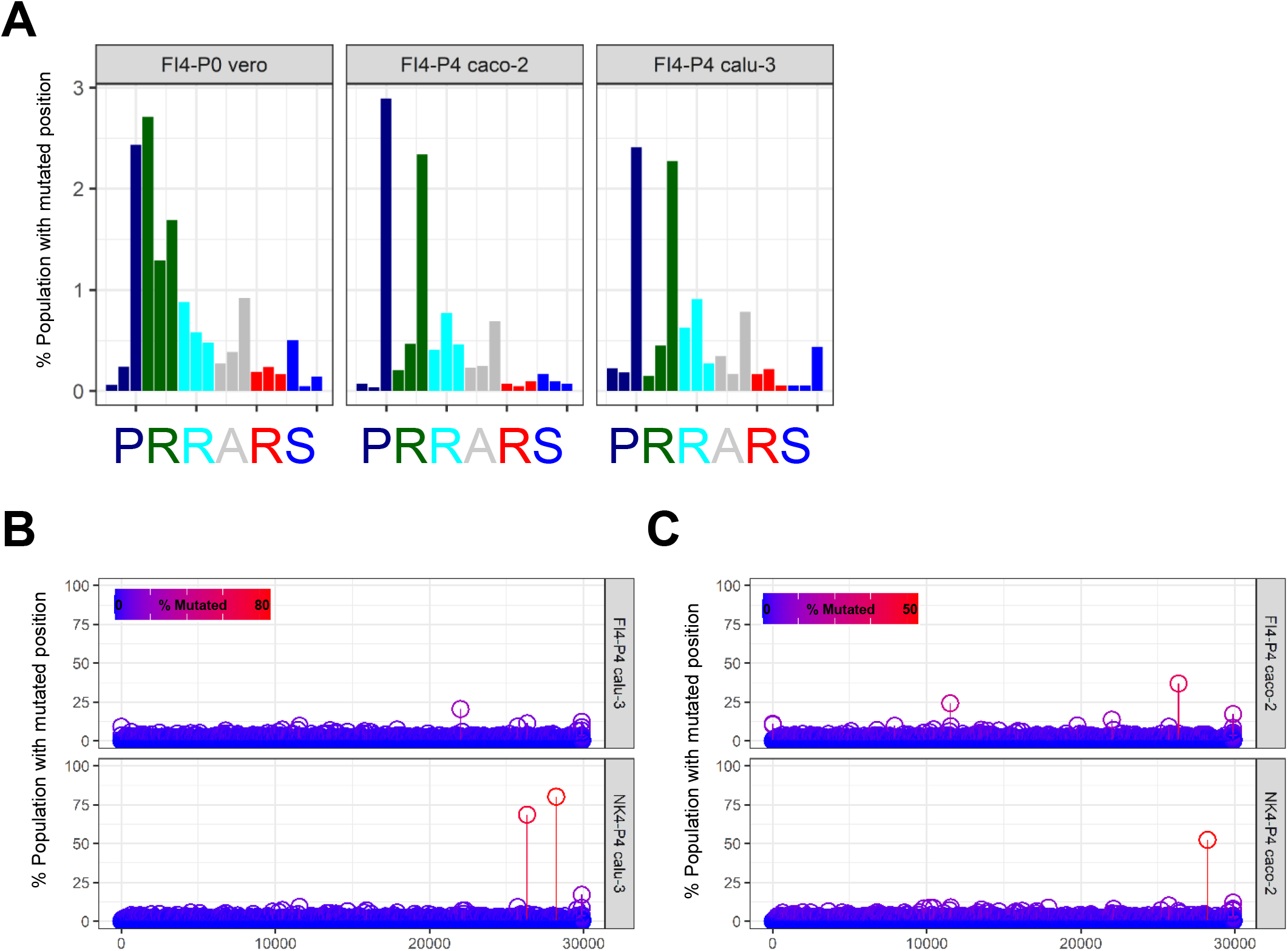

**Figure S4.**
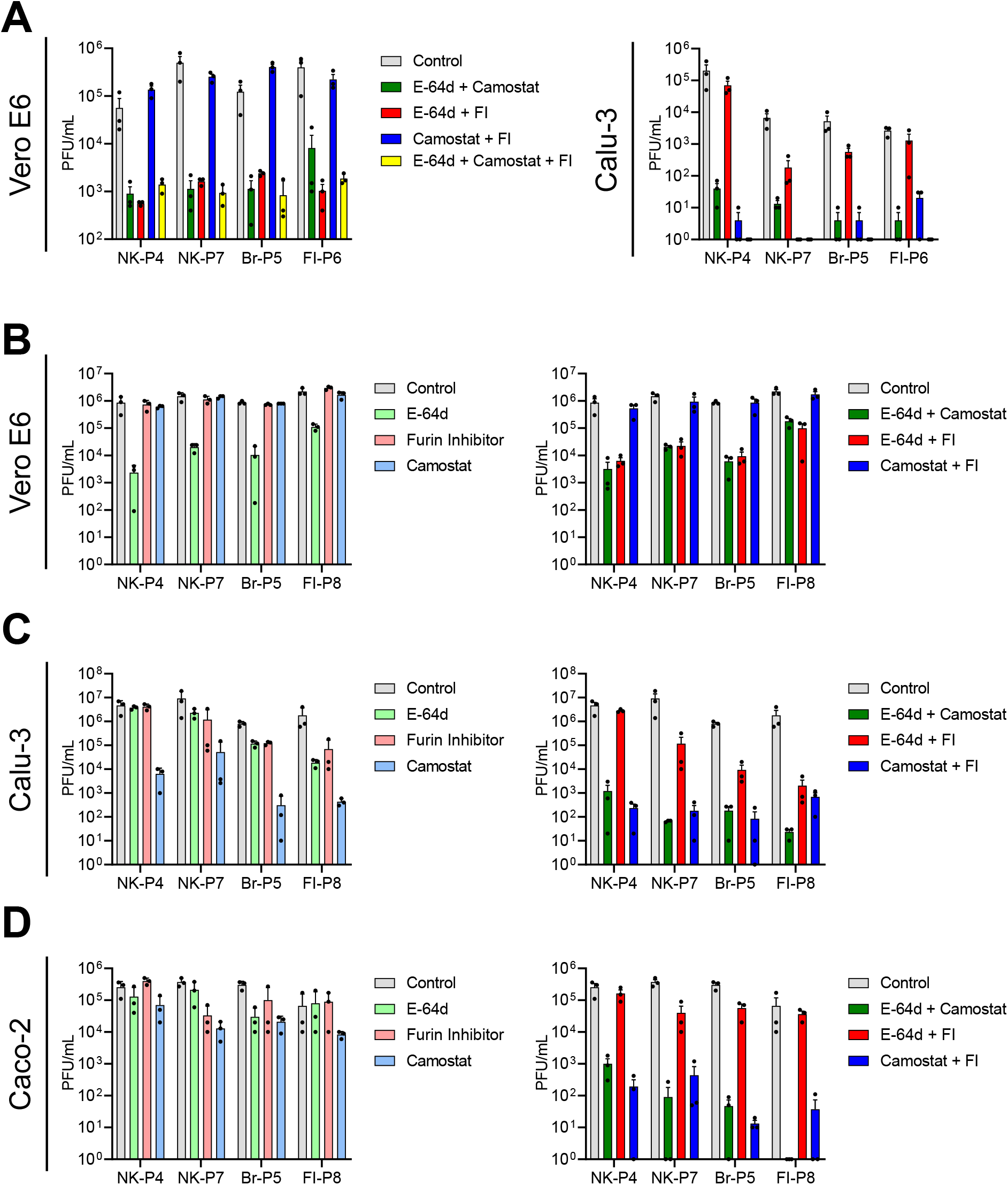

**Figure S5.**
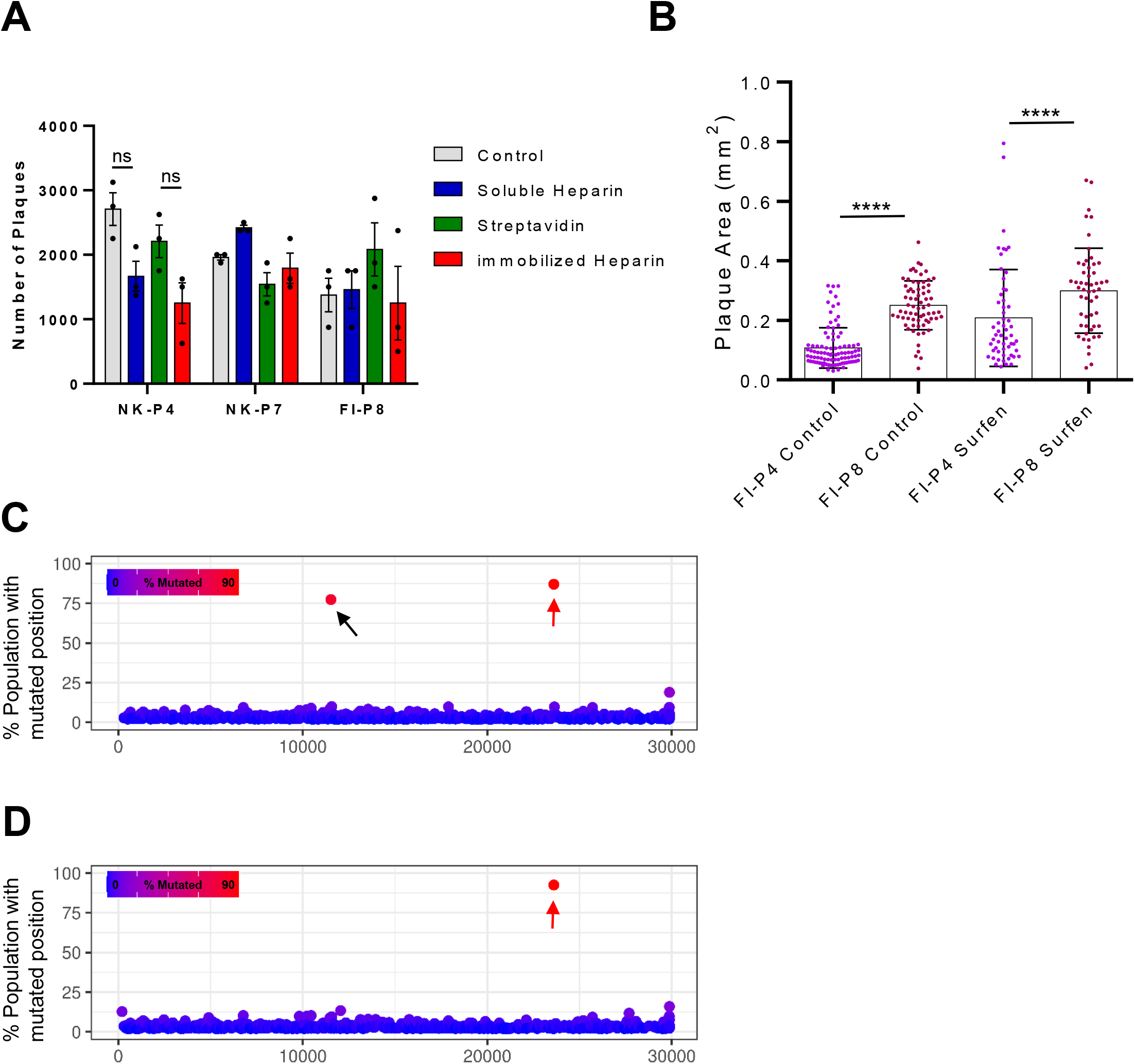

