## Supplemental Data for "Rapid SARS-CoV-2 Adaptation to Available Cellular Proteases"

#### Figure S1

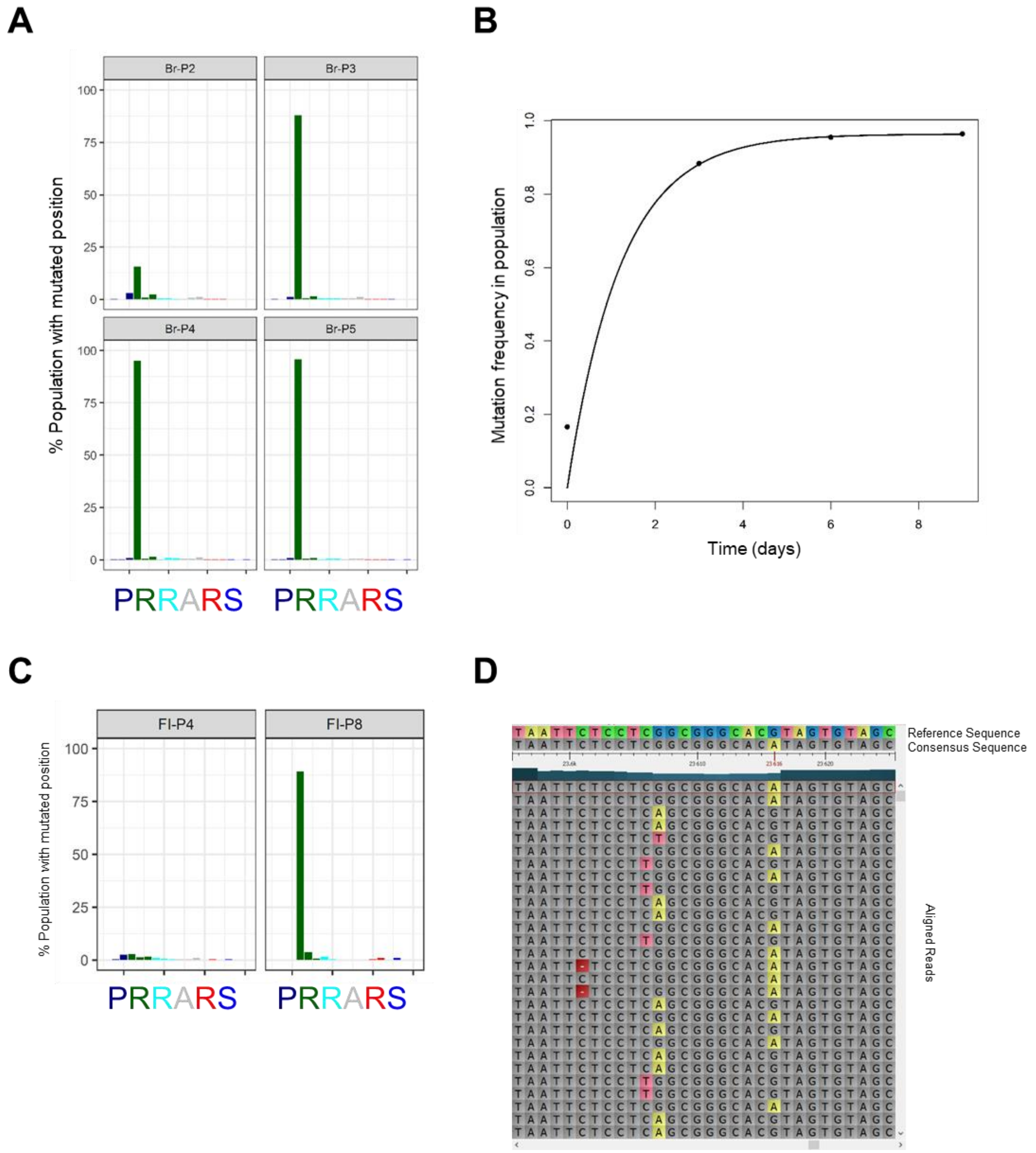


**Figure S1. Multiple strains of SARS-CoV-2 quasispecies show mutation at the furin cleavage site after serial passages on Vero E6, Related to Figure 1.**

(A) The Br strain was serially passaged on Vero E6 cells (Br-P2 until P5). Mutation frequencies of nucleotides at the furin cleavage site in SARS-CoV-2 S are plotted on y-axis. Three bars of same color represent the nucleotide codon triplet for each corresponding amino acid at the furin cleavage site from genomic position 23,603 to 23,620 (x-axis).

(B) The sum of mutation frequency at the furin site position 23,606, 23,607 and 23,616 of Br strain serially passaged on Vero E6 cells is plotted against time.

(B) The sum of mutation frequency at furin site position 23,606, 23,607 and 23,616 of Br strain serially passaged on Vero E6 cells is plotted against time. The mutation rate and reverse mutation rate correspond to change per day calculated by mathematical model.

(C) The FI strain was serially passaged on Vero E6 cells. The mutation frequency of nucleotides at furin cleavage site in SARS-CoV-2 S for passage 4 and 6 (FI-P4 and FI-P8) are plotted on y-axis. Three bars of same color represent the nucleotide codon triplet for each corresponding amino acid at the furin cleavage site from genomic position 23,603 to 23,620 (x-axis).

D) The NK strain passage 6 (NK-P6) alignment file (.bam) is aligned to the reference genome with the help of Tablet. Mutated positions were highlighted in color in otherwise grey consensus sequence and individual reads. The visual analysis was performed to check if position 23,606, 23,607 and 23,616 mutations were present on same reads, and it was observed that they were present on mutually exclusive reads.

The genomic positions and reference file used correspond to Wuhan-Hu-1 isolate (GenBank accession no: NC_045512).

#### Figure S2

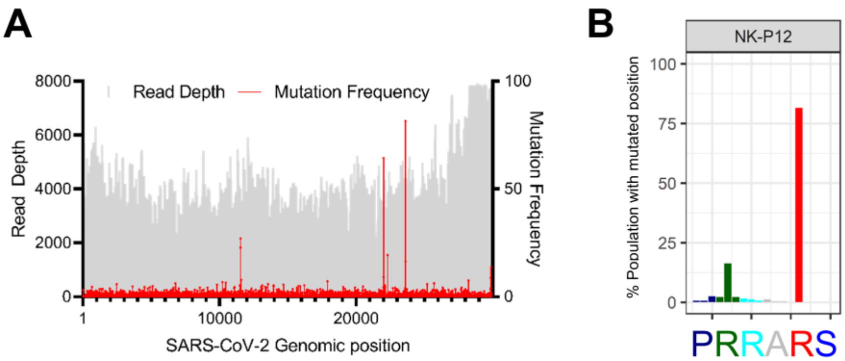


**Figure S2. The highly passaged NK-P12 strain maintains wildtype furin cleavage site in viral quasispecies, Related to Figure 1.**

The NK strain was passaged 10 times on Vero E6 cells and NK-P12 (passage 12), and analyzed with deep sequencing to access mutation in the virus genome.

(A) Read Depth (grey bar) and mutation frequency (red dots) for whole NK-P12 genome are plotted. (B) The mutation frequency of nucleotides at the furin cleavage site are plotted on y-axis. Three bars of same color represent the nucleotide codon triplet for each corresponding amino acid at the furin cleavage site from genomic position 23,603 to 23,620 (x-axis). The genomic positions correspond to Wuhan-Hu-1 isolate (GenBank accession no: NC_045512).

### Figure S3

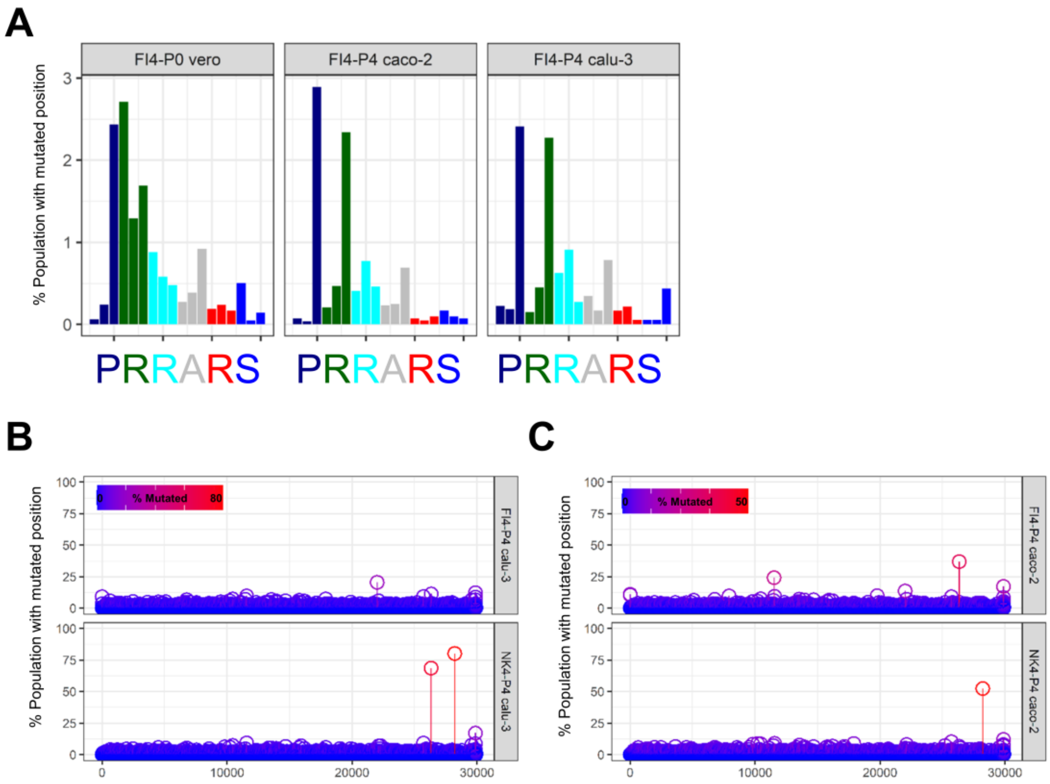


**Figure S3. SARS-CoV-2 quasispecies composition of different stains at furin cleavage after passages on TMPRSS2 competent cell lines, Related to Figure 3.**

(A) The FI stain, passaged 4 time initially on Vero E6 cells (FI-P4), was serially passaged on Calu-3 or Caco-2 cells. The three graphs juxtapose the mutation frequency at the furin cleavage site of the viral population directly after passing on Vero E6 (FI4-P0 vero) to populations that were further grown on either Calu-3 (FI4-P4 calu-3) or Caco-2 (FI4-P4 caco-2). The mutation frequency of nucleotides at the furin cleavage site are plotted on y-axis. Three bars of same color represent the nucleotide codon triplet for each corresponding amino acid at the furin cleavage site from genomic position 23,603 to 23,620 (x-axis).

(B-C) NK and FI strain initially passaged 4 times on Vero E6 cells were further passaged on Calu-3 (B) and Caco-2 (C). Mutation frequency for whole genome is plotted on y-axis against genomic position (x-axis). Increased mutation frequency is highlighted by symbol color shift from blue to red.

Figure S4

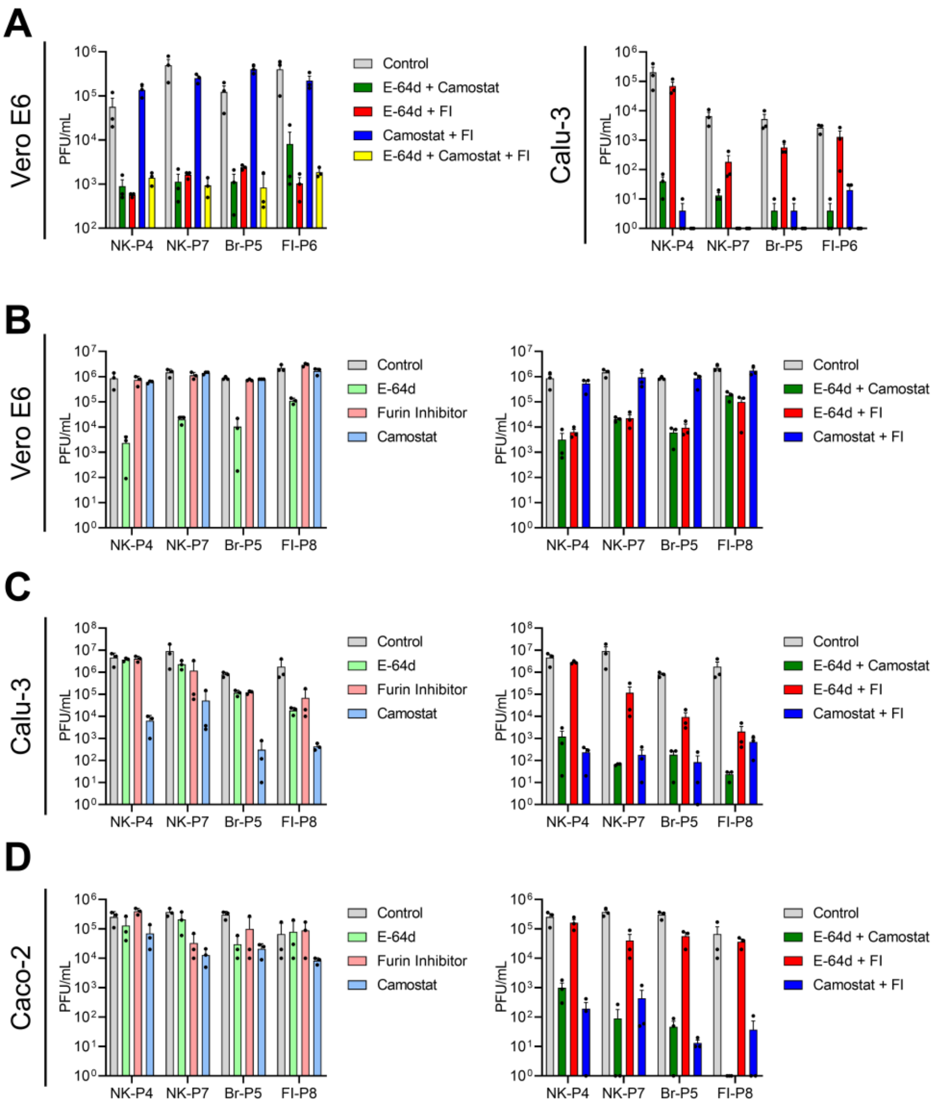


**Figure S4. SARS-CoV-2 growth inhibition at day 1 and 2 post infection by different protease inhibitors, Related to Figure 4.**

(A) Vero E6 and Calu-3 cells were infected at an MOI of 0.01 in the presence or absence of different protease inhibitor combinations. The supernatant from infected cells was collected at 24 hpi and titrated on Vero E6 cells.

(B-D) Vero E6 (B), Calu-3 (C) and Caco-2 (D) cells were infected at an MOI of 0.01 in the presence or absence of different protease inhibitors. The supernatant from infected cells was collected at 48 hpi and titrated on Vero E6 cells. Error bars represent mean ± SEM of three biological replicates.

Figure S5

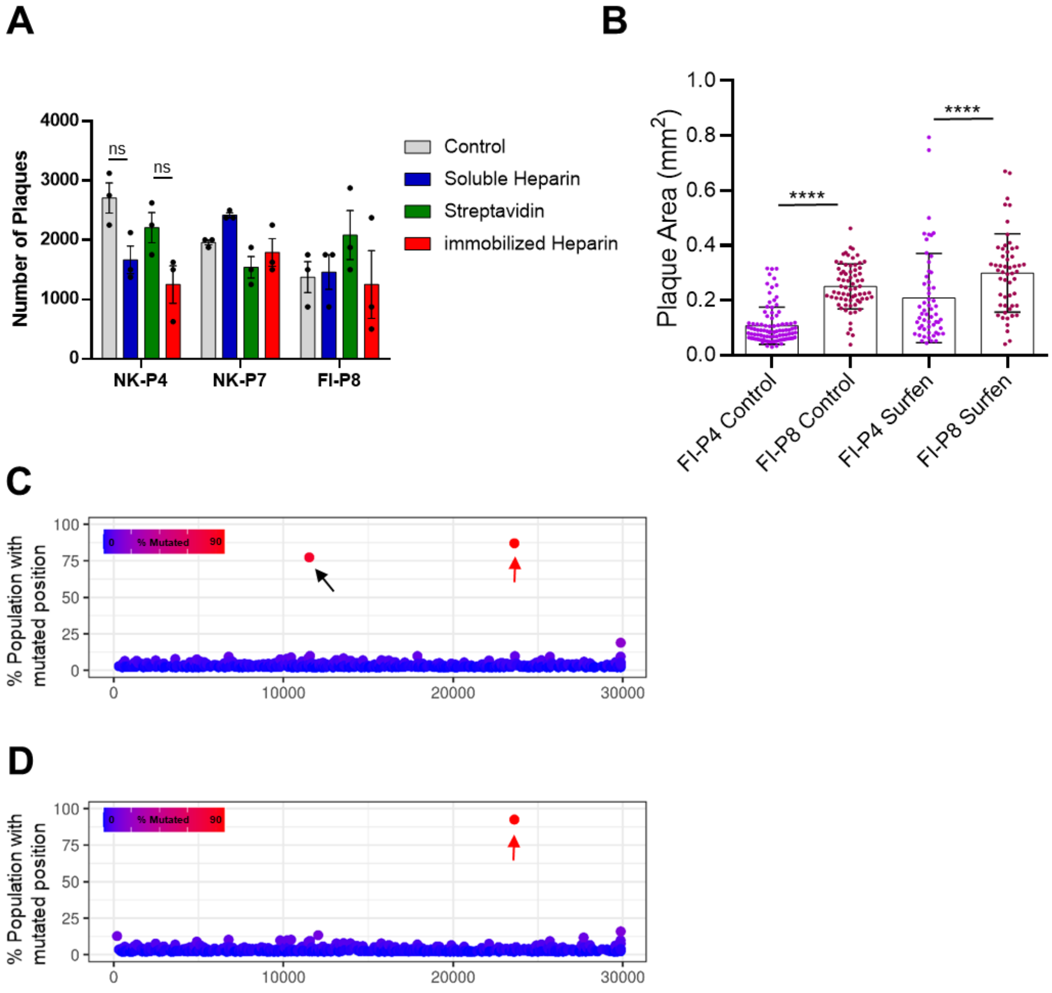


**Figure S5. SARS-CoV-2 cell entry and growth is not affected by heparin pre-treatment or HS antagonists.**

(A) Heparin binding was tested by incubating NK-P4, NK-P7 and FI-P8 virus suspension with Heparin salt or Biotin-Heparin immobilized on Streptavidin coated plate. After one-hour incubation at 37°C, virus was titrated on Vero E6 cells.

(B) Vero E6 cells were infected with low or high-passage FI strain. The cells were overlaid with methylcellulose supplemented with 10 µM surfen. The virus plaque size was quantified 3 dpi. Each symbol represents one plaque and data is pooled from multiple infected wells of two independent experiments. Statistical significance was calculated using one way ANOVA and Bonferroni posttest. ****p > 0.0001.

FI (C) and Br (D) strains were passaged in Vero E6 cells in the presence of 10 µM surfen. After 4 passages, virus genome sequence was analyzed with deep sequencing. Each symbol represents an individual nucleotide, and genomic positions (x-axis) with mutation frequency >1% are plotted. Red arrows highlight the position of the furin cleavage site and black arrow show mutation in nsp6 resulting in F3753V.

**Table S1. Mutations in SARS-CoV-2 E protein upon passage in different cell lines, Related to Figure 3.**

| SARS-CoV-2 Strain | Cell line & Passage #[a] | Mutation (position)[b] Nucleotide[c] | Amino Acid[d] | Mutated Population |
|---|---|---|---|---|
| Sudtirol-FI4[e] | P0 | C->T (26333) | T->I (30) | 4.30% |
| Sudtirol-FI4 | Vero P4 | C->T (26333) | T->I (30) | <1% |
| Sudtirol-FI4 | Caco-2 P4 | C->T (26333) | T->I (30) | 36.83% |
| Sudtirol-FI4 | Calu-3 P4 | C->T (26333) | T->I (30) | 11.28% |
| Ischgl-NK4[f] | P0 | T->C (26324) | L->S (27) | 9.7% |
| | | C->T (26333) | T->I (30) | <1% |
| Ischgl-NK4 | Vero P4 | T->C (26324) | L->S (27) | <1% |
| | | C->T (26333) | T->I (30) | <1% |
| Ischgl-NK4 | Caco-2 P4 | T->C (26324) | L->S (27) | 6.49% |
| | | C->T (26333) | T->I (30) | 1.97% |
| Ischgl-NK4 | Calu-3 P4 | T->C (26324) | L->S (27) | 68.58% |
| | | C->T (26333) | T->I (30) | <1% |
| Ischgl-NK6[g] | P0 | T->C (26324) | L->S (27) | <1% |
| | | C->T (26333) | T->I (30) | <1% |
| Ischgl-NK6 | Caco-2 P4 | T->C (26324) | L->S (27) | 61.62% |
| | | C->T (26333) | T->I (30) | 1.98% |
| Ischgl-NK6 | Calu-3 P4 | T->C (26324) | L->S (27) | 1.37% |
| | | C->T (26333) | T->I (30) | 4.67% |

[a] Passage number on the given cell line.
[b] Positions with Mutation frequency above 1% are shown.
[c] Numbers are according to Wuhan-Hu-1 sequence (GenBank accession no: NC_045512).
[d] Numbers start from the amino terminus of protein E.
[e] FI4 represents input virus, FI strain passaged 4 times on Vero E6 before passage on a given cell line. FI4 strain did not show positon 26324 mutation above 1% in any cell line.
[f] NK4 represents input virus, NK strain passaged 4 times on Vero E6 before passage on a given cell line.
[g] NK6 represents input virus, NK strain passaged 6 times on Vero E6 before passage on a given cell line.

#### Data S1. R code used for mathematical modelling.

```
######################################################
### Fits a curve of the form y(t) with y(0)=0 as
### described in the paper to given data points.
### tps: Time points
### var.freqs: Observed mutation frequencies
### err.func: Error function. Here the sum of
###         absolute errors is used (instead
###         of the sum of squared errors
###         which is more sensitive outliers.)
### Value:
```

```
### A vector of two fitted parameters corresponding
### to a and (a+b) in the paper.
##########################################################
fit.mut.curve <- function(tps,var.freqs,err.func=abs){
  ## Error function
  obj.func <- function(params){
    ## Mutation rates must be between 0 and 1.
  if (params[1]<0 | params[1]>1 | (params[2]-params[1])<0 | (params[2]-params[1])>1)
                    {return(.Machine$double.xmax)}
                   ## Te parameter for a+b must not be smaller than the one for a.
                   if (params[2]<params[1]){return(.Machine$double.xmax)}
  return(sum(err.func((params[1]/params[2])*(1-exp(-params[2]*tps)) - var.freqs)))
  }
##Fit the parameters with optim.
return(optim(c(0.5,0.5),obj.func,method="SANN")$par)
}
#The indices of the considered positions for the mutation (positions 23606, 23607, 23616).
mutation.positions <- c(4,5,14)
## Read the data for the calu strain. The directory path might need adjustment.
datfurin <- read.table(file="C:/Users/zch14/Desktop/cov2/Alternate frequency/calu_NK4_furin_detail.tab",header=T,sep="\t")

##Time points for the calu strain.
tps <- 3*(0:4)

###Compute the mutation frequencies for the ime points.
varfr <- rep(0,5)
for (i in 1:length(varfr)){
  varfr[i] <- sum(datfurin[(i-1)*18+ mutation.positions,3])/100
}
##Fit the curve.
sa <- fit.rev.mut.curve(tps,varfr)

##Plot the curve and the points.
plot(tps,varfr,pch=16,xlab="t[days]",ylab="Mutation frequency",ylim=c(0,max(varfr)))
curve((sa[2]-sa[1])/sa[2]*exp(-sa[2]*x) +sa[1]/sa[2],add=T,lwd=2)
text(6,0.6,paste0("Mutation rate=",round(sa[1],2),", Reverse mutation rate=",round(sa[2]-sa[1],2)))
##################################################################################
```
